# Development of a PNGase Rc column for online deglycosylation of complex glycoproteins during HDX-MS

**DOI:** 10.1101/2023.07.28.550801

**Authors:** Thomas Lambert, Marius Gramlich, Luisa Stutzke, Luke Smith, Dingyu Deng, Philipp D. Kaiser, Ulrich Rothbauer, Justin L.P. Benesch, Cornelia Wagner, Maximiliane Koenig, Petr Pompach, Petr Novak, Anne Zeck, Kasper Rand

## Abstract

Protein glycosylation is one of the most common PTMs and many cell surface receptors, extracellular proteins and biopharmaceuticals are glycosylated. However, HDX-MS analysis of such important glycoproteins has so far been limited by difficulties in determining the HDX of the protein segments that contain glycans. We have developed a column containing immobilized PNGase Rc (from *Rudaea cellulosilytica*) that can readily be implemented into a conventional HDX-MS setup to allow improved analysis of glycoproteins. We show that HDX-MS with the PNGase Rc column enables efficient online removal of N-linked glycans and the determination of the HDX of glycosylated regions in several complex glycoproteins. Additionally, we use the PNGase Rc column to perform a comprehensive HDX-MS mapping of the binding epitope of a mAb to c-Met, a complex glycoprotein drug target. Importantly, the column retains high activity in the presence of common quench-buffer additives like TCEP and urea and performed consistent across 114 days of extensive use. Overall, our work shows that HDX-MS with the integrated PNGase Rc column can enable fast and efficient online deglycosylation at harsh quench conditions to provide comprehensive analysis of complex glycoproteins.

## INTRODUCTION

Glycosylation of Asn residues (N-glycosylation) is a predominant post-translation modification of proteins exposed to the extracellular environment^1,2^. N-glycosylation offers structural stability, improved solubility, and binding partner recognition mechanisms that allow proteins to carry out their function outside of cells^3^. Also, many extracellular proteins of interest as targets for drugs or as drug modalities (biopharmaceuticals) are N-glycosylated^1,2^. For these reasons, there is a growing need to be able to study the structure and interactions of proteins in their natively glycosylated state^4^. However, traditional methods for structural analysis of proteins like x-ray crystallography and NMR are often not readily compatible with natively glycosylated proteins due to the increased dynamics and structural heterogeneity afforded by the presence of N-glycosylations.

Hydrogen-deuterium exchange mass spectrometry (HDX-MS) can provide a powerful alternative method to study the conformation and interactions of dynamic or conformationally-heterogeneous protein systems in solution^5^. In HDX-MS, a protein is subjected to deuterated solvent and the exchange of the backbone amide hydrogens for heavier deuterium atoms of the solvent can be measured by MS. Involvement in hydrogen-bond networks, which define protein higher-order structure, heavily influences the exchange rate of backbone amide hydrogens, making HDX-MS a sensitive probe of protein conformation and dynamics. The HDX of proteins can be resolved to the peptide-level using online proteolysis. The sensitivity of HDX-MS and its intrinsic ability to tolerate sample complexity makes the method helpful for studying a variety of protein states, like glycoproteins, that can be incompatible to conventional methods^5^. Common applications of HDX-MS include analysis of conformational changes in proteins during function and regulation, including those of membrane proteins, protein-ligand/drug interactions, mode of action of biopharmaceuticals and epitope mapping of therapeutic monoclonal antibodies^5–8^.

Glycosylation can however present particular difficulties for HDX-MS. Different glycoforms of a protein can coexist when various heterogenous glycan structures occupy a single site^9,10^ and as a result cause glycan-induced signal spread, which results in a weaker signal during MS analysis and thus gaps in the sequence coverage in regions containing N-linked glycosites. Additionally, glycan HDX interpretation may be impacted by the retention of deuterium by acetamido groups within glycan structures which introduces error when quantifying deuterium retention^9,11^.

One solution is to use fragmentation in MS/MS analysis as a means of overcoming these difficulties^12,13^. However, collision-induced dissociation (CID) MS/MS is especially difficult to use for this purpose. This is due to the preferential fragmentation of glycosidic links in glycopeptides^12,13^ and the occurrence of H/D scrambling during CID^14^. Other fragmentation methods that occur without scrambling, such electron transfer dissociation (ETD)^15^, have demonstrated selective cleavage of the peptide backbone-while keeping the glycosidic bond intact. However, these options require MS systems having this capability and ETD suffers from markedly lower fragmentation efficiency than CID^12,13^.

N-glycan removal during HDX-MS have also been explored using acid-stable endoglycosidases. The first enzyme to be used for this purpose was PNGase A, which was used in an off-line manner to facilitate deglycosylation at quench conditions during HDX-MS of glycoproteins^9^. However, the relatively high pH optimum of 5 of PNGase A and its low tolerance to common quench buffer additives like TCEP and urea, prompted the use of other acidic PNGases, like PNGase H+, (from *Terriglobus roseus*)^10^. Similar to PNGase A, PNGase H+ can release a broad variety of N-glycans and has the advantage over PNGase A, that it can also deglycosylate glycoproteins and not just glycopeptides^9^. Another advantage over PNGase A is its pH optimum at 2.6 and high tolerance to reducing agents like TCEP. However, PNGase H+ showed low expression yields and purity. Later on, a few PNGase H+ variants were found and some of them showed high deglycosylation activities. One of the promising PNGase H+ variants was PNGase Dj originating from *Dyella japonica*. To facilitate the use of such rare enzymes we immobilized PNGase Dj to a microchip and showed its use in HDX-MS during analysis of complex glycoproteins including epitope mapping applications^16^. Building on this work, we have recently shown that another PNGase H+ variant, PNGase Rc originating from *Rudaea cellulosilytica*, has higher intrinsic activity and higher expression yields than Dj, the latter likely due to the presence of only two disulfide bonds in PNGase Rc vs three in PNGase Dj. Furthermore, PNGase Rc can cleave a broad variety of N-glycans (similar to PNGase H+), including core-fucosylated glycan structures (not cleaved by PNGase F) and glycans from plants and invertebrates^17^. For these reasons, PNGase Rc represents an attractive enzyme for use in HDX-MS applications of glycoproteins^18,19^.

Here we have developed a column containing immobilized PNGase Rc that can be used in an online HDX-MS workflow to greatly improve analysis of glycoproteins. The immobilization of PNGase Rc offers considerable advantage compared to the usage of PNGase enzymes in solution, including reusage and a higher enzyme to substrate ratio can be achieved. Additionally, of key advantage to HDX-MS, no pre-incubation time is needed, resulting in a fast online workflow that increases throughput and limits back-exchange^16^. We show that the PNGase Rc column enables efficient online removal of N-linked glycans and the determination of the HDX of glycosylated regions in several highly complex/challenging glycoproteins, including in epitope mapping applications. The column retains activity in the presence of common HDX-MS quench-buffer additives like TCEP and urea and maintains high deglycosylation activity after repeated use across >100 days.

## EXPERIMENTAL SECTION

### PNGase Rc Expression and Purification

See Supporting information and as described previously^18,19^

### Immobilization of PNGase Rc and column packing

See the Supporting information. On-column immobilization of PNGase Rc was performed by AffiPro Ltd.

### HDX-MS of glycoproteins using the PNGase Rc column

Haptoglobin and transferrin experiments were performed using a Synapt G2Si HDMS coupled to an HDX-MS UPLC system (Waters Corporation, UK).

HDX of Haptoglobin (Hp 1-1, Sigma-Aldrich) was initiated by diluting the protein into labelling buffer (20 mM Tris pD 7.0, 90% D_2_O) to a concentration of 1 μM. The HDX reaction was quenched at various timepoints by transferring reaction aliquots (50μL)) into a 1:1 (v/v) volume of quench buffer (250mM TCEP and 4M Urea in 0.5M Glycine at pH 2.3). Quenched samples were incubated on ice for 3 minutes to allow for protein unfolding and reduction and frozen on dry-ice. During analysis, HDX samples were rapidly thawed and injected into the cooled (0°C) HDX-MS UPLC system and subjected to online proteolysis (20°C) in an in-house built column packed with agarose-immobilized pepsin, followed by online deglycosylation (20°C) using a column placed immediately downstream packed with immobilized PNGase Rc (2.1 × 20 mm, Wuxi GALAK Chromatography Technology Co, China), and finally desalting on a trap column (Waters Acquity BEH C18, 1.7 μm, 2.1 mm x 5 mm length) for 3 min with aqueous phase (0.23% FA) at 200μl/min. Separation was performed on an analytical column (Waters Acquity BEH C18, 1.7 μm, 1 mm × 100 mm) using a 9 min gradient from 2 to 40% organic phase (0.23% FA in acetonitrile) at 0°C. Peptides were identified by both data-dependent acquisition (DDA) and data-independent acquisition (DIA) mode (i.e. MSe) MS/MS and were filtered using Protein Global Server 3.0 (Waters) and DynamX v3.0 (Waters). The deuterium uptake of respective peptides was determined using DynamX. The HDX experiment was performed in independent triplicates.

Maximum-labeled control samples of Hp 1-1 were generated by online digestion of Hp 1-1 as described above and manual collection of samples after trapping. Peptic peptides of Hp 1-1 were exposed to labelling buffer as above for 5 hrs at to achieve maximal labelling and applied to the HDX-UPLC system configured and operated exactly like described above, but without the pepsin column and in both the absence and presence of the PNGase Rc column.

Back-exchange (BE) values were calculated by comparing the data from the maximal deuterated control sample against an undeuterated control.

HDX of transferrin was performed by diluting solid apo-human transferrin (Sigma-Aldrich to a concentration of 10 μM in equilibration buffer (PBS, pH 7.4). Labelling was initiated by diluting 5 μL of each protein aliquot to 50 μL total volume by addition of labelling buffer (PBS prepared in D_2_O, pD 7.0) giving a 90% D_2_O environment. Labelling took place for various time points. The labelling reaction was quenched by a 1:1 addition of ice-cold quench buffer (50mM KH_2_PO_4_, 4M Urea, 200mM TCEP) to a final pH of 2.5. Samples were digested online (20°C) with a Waters Enzymate BEH pepsin column, followed by online deglycosylation (20°C) using a column placed immediately downstream packed with immobilized PNGase Rc (2.1 × 20 mm, Wuxi GALAK Chromatography Technology Co, China). Peptides were trapped on a Waters BEH C18 VanGuard pre-column for 3 min at a flow rate of 100 μL/min in buffer A (0.1% formic acid ~ pH 2.5).

Peptides were separated on a Waters BEH C-18 analytical column with a linear gradient of buffer B (0.1% formic acid in acetonitrile ~ pH 2.5) at a flow rate of 40 μL/min. All trapping and separation were performed at ~0°C to reduce back exchange. All data was collected using the same flow rates for states with and without the PNGase column.

MS data was acquired using MSE workflow. All time points across this experiment were obtained in triplicate. MS was calibrated separately using NaI and the MS data was obtained with lock mass correction using LeuEnk. Peptides were assigned with the ProteinLynx Global Server (PLGS, Waters Corporation, UK) software and the isotope uptake of respective peptides was determined using DynamX v3.0.

HDX of VISTA was performed by reconstituting the protein (Human B7-H5, Acro BioSystems) 5x concentrated compared to manufacturer’s instructions. The reconstituted solution contained 50 % trehalose which inhibited in pre-experiments deglycosylation by PNGase Rc. Therefore, VISTA was buffer-exchanged using Zeba Spin Desalting Columns (7K MWCO Fisher Scientific, USA) to PBS buffer. The final concentration determined via NanoDrop 2000 spectra photometer (Fisher Scientific, USA) was 1.65 mg/mL.

6 μL VISTA (90 μM in PBS) were deuterated by incubation with 54 μL PBS HDX buffer (1x, D_2_O pH 7.4) for 0.25, 1, 10, 60 and 600 min at 25°C. Aliquots of 10 μL were quenched and reduced by adding 1:1 (v/v) ice-cold quenching solution (0.4M TCEP and 4M urea in 100mM ammonium formate, pH 2.4) resulting in a final pH of 2.6 and a final concentration of 0.2 M TCEP and 2M urea. Samples were immediately snap-frozen in liquid nitrogen and stored until analysis at -80°C. All experiments were performed in independent triplicates.

Full deuteration labelling was attempted by deuteration of the above-described solution for 50 h and by incubation of VISTA in the presence of 2M urea-d4 prepared in D2O. Samples were incubated at 20°C and quenched by adding 1:1 (v/v) ice-cold quenching solution (0.4M TCEP dissolved in 100mM ammonium formate buffer, pH 2.6). Full deuteration was however not achieved under the tested conditions.

### HDX-MS epitope mapping of c-Met

C-Met (Proteros Biostructures GmbH, Germany) experiments were performed using an Orbitrap Fusion™ Lumos™ (Thermo Fisher Scientific, US) coupled to an Acquity UPLC system with HDX (Waters corporation, UK. Sample preparation was performed by the LEAP HDX autosampler (Trajan).

HDX of unbound and bound c-Met (1:2.2 ratio with mAb (Roche Diagnostics GmbH, Germany)) was performed by diluting c-Met into labelling buffer (20mM Histidine, 140μM NaCl, 90% D_2_O, pD 6.0) to a concentration of 0.35μM. The labelling reaction (60μl) was quenched at various time-intervals by a 1:1 addition of ice-cold quench buffer (170mM KH_2_PO_4_, 250mM TCEP 4M Urea, pH 2.8). HDX samples were injected into the cooled (0°C) HDX-MS UPLC system and subjected to online proteolysis (20°C) with a Nepenthesin-2 column (AffiPro), followed by online deglycosylation (20°C) using a column placed immediately downstream packed with immobilized PNGase Rc and trapped and desalted on a Waters BEH C18 VanGuard pre-column for 3 min at a flow rate of 200 μL/min in buffer A (0.23% formic acid ~ pH 2.5). Peptides were separated on a Waters BEH C-18 analytical column using a linear gradient of buffer B (0.23% formic acid in acetonitrile ~ pH 2.5) at a flow rate of 40 μL/min. All trapping and separation were performed at ~0°C to reduce back exchange. All data was collected using the same flow rates for states with and without the PNGase Rc enzymatic column.

Peptide identification was performed in three runs by DDA fragmentation using HCD, ETD and EThcD and the following criteria in the PEAKS Studio 11 (Bioinfomatic Solutions, inc): Peptide tolerance: 5 ppm, MS/MS tolerance: 0.5 Da. The deuterium uptake of respective peptides was determined using HDExaminer 3.0 (Sierra Analytics).

### Tryptic digestion of Trastuzumab (TZ)

See the Supporting Information and as described previously^16^

### LC-MS assay for monitoring the online deglycosylation activity of the PNGase Rc column

Assessment of PNGase Rc activity was done in a similar manner as described previously using samples from a tryptic digest of Trastuzumab (TZ)^16^. The PNGase Rc column was interfaced with a cooled HDX-UPLC system (Waters, Milford, MA). The LC-MS configuration and columns used were the same as during HDX-MS experiments of Hp 1-1. 50pmol of TZ tryptic digest was injected and run through the column at 200μl/min of LC mobile phase (0.23% formic acid (FA), pH

2.5) before desalting and separation by reversed-phase chromatography and MS analysis. As described previously^16^, the online deglycosylation activity of the column was evaluated by monitoring the signal intensity of the glycosylated and deglycosylated tryptic peptide EEQYNSTYR normalized to the signal intensity of the ATII peptide internal standard.

## RESULTS AND DISCUSSION

### Immobilization of PNGase Rc

PNGase Rc was recombinantly expressed in *E*.*coli* as described previously at high yield and purity^18,19^. Immobilization was performed on POROS™ AL beads via Lys residues in PNGase Rc. Investigation of the AlphaFold-generated structure of PNGase Rc revealed that all Lys residues were located outside of the predicted active site (based on looking at active sites of PNGase F and A, Figure S1)^18^. The immobilization efficiency (Ψ) was tested by assessment of PNGase Rc concentration in the supernatant after bead coupling. The concentration was determined from a calibration curve using the UV 280nm peak area of standard dilutions of PNGase Rc analyzed by HPLC using a C4 column (Figure S2). An enzyme-to-bead (E:B) ratio of 1:10 yielded ~ 39% immobilization efficiency, whereas an E:B ratio of 1:20 yielded 95% immobilization efficiency. Therefore, the E:B ratio of 1:20 results in quantitative coupling of the enzyme to the beads but most likely not the whole accessible surface is modified with enzyme. The activity of immobilized PNGase Rc was assessed as described previously using an LC-MS assay monitoring the deglycosylation of the tryptic glycopeptide encompassing the N297 glycosite of the IgG1 Trastuzumab (TZ)^9,16,19^. Immobilization with 1:10 E:B resulted in beads with significantly higher deglycosylation efficiency than beads produced by immobilization with 1:20 E:B (or PNGase Rc free in solution) (Figure S4). Thus, beads made using 1:10 E:B ratio were selected for further study and column packing.

Haptoglobin 1-1 (Hp 1-1) was chosen as the model substrate to further assess the deglycosylation activity of bead-immobilized PNGase Rc. Hp 1-1 is one of the three phenotypes (Hp 1-1, Hp 2-1, Hp 2-2) of the human glycoprotein haptoglobin. Hp 1-1 is composed of a light α-chain and a heavy β-chain (Figure 1A) and forms a disulfide-bonded dimer^20^. Four glycans are attached to the β-chain and are located at residues N23, N46, N50, and N80.

**Figure 1:**
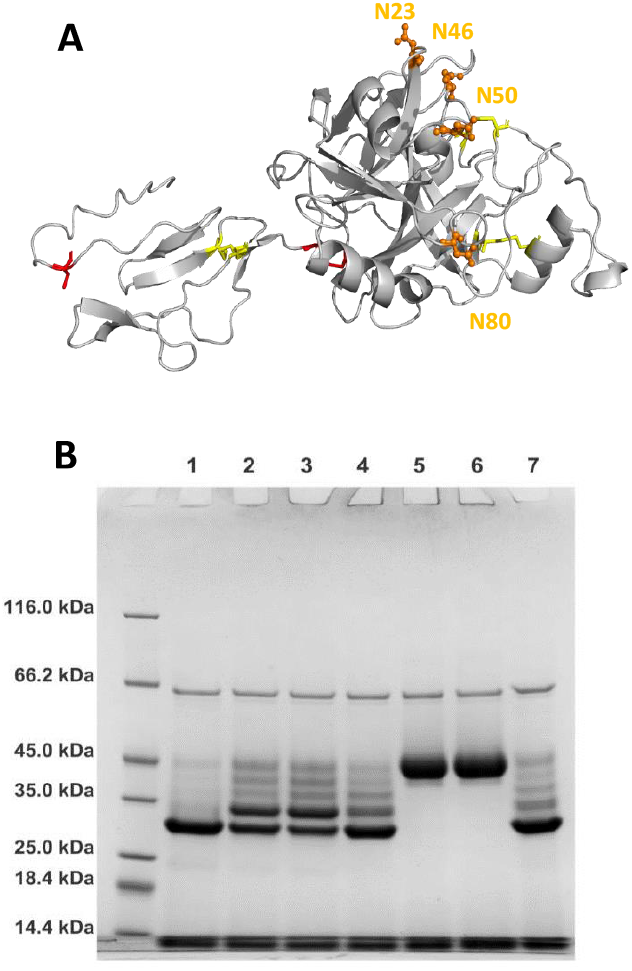
A) Structure of the Hp 1-1 monomer (AlphaFold) containing both the alpha and beta chain. Disulfide bonds are shown in yellow (intra) and red (inter) sticks, while glycans are shown in orange sticks and labeled. B) Activity of immobilized PNGase Rc displaying the appearance of bands corresponding to deglycosylated species of Hp 1-1 upon treatment with PNGase Rc at various pH values. In all cases, the same amount of Hp 1-1 and PNGase Rc was used. L1-L6: pH 2.4, pH 3.4, pH 4.4, pH 5.4, pH 6.4, pH 7.4 and L7:0.1% formic acid. Glycosylated Hp 1-1 beta chain (∼45 kDa) was converted into bands corresponding to removal of 1, 2, 3 and all 4 N-glycans (∼30 kDa for fully deglycosylated form).

PNGase Rc immobilized on beads was capable of full deglycosylation of all N-linked glycans of Hp 1-1 at pH 2.4 as monitored by SDS-PAGE (Figure 1B). Upon deglycosylation, the band corresponding to glycosylated Hp 1-1 beta chain (∼45 kDa) was converted into bands corresponding to removal of 1, 2, 3 and all 4 N-glycans (∼30 kDa for fully deglycosylated form). At higher pH values the reaction was less efficient, in good agreement with the reported pH optimum of PNGase Rc^18,21^. The PNGase Rc-functionalized beads (1:10 E:B ratio) were packed into a HPLC/UPLC compatible column (2.1 × 20 mm) and integrated into a conventional HDX-MS setup (Figure 2).

**Figure 2:**
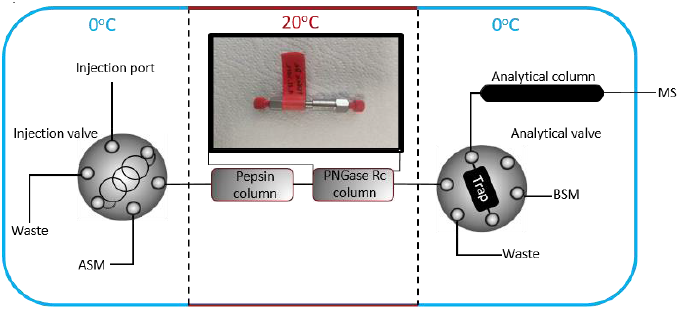
Integration of the PNGase Rc column within a commercially available HDX-MS setup.

### HDX-MS of glycoproteins of varying complexity

To evaluate the implementation of the PNGase Rc column into HDX-MS setups in different labs, PNGase Rc columns were sent to three separate research labs and were used to perform HDX-MS experiments on three different glycoproteins of significant complexity in terms of disulfide-bonds and N-linked glycans. Namely, Hp 1-1, V-domain Ig suppressor of T cell activation transmembrane protein (VISTA) and transferrin..

As mentioned previously, Hp 1-1 forms a disulfide-bonded dimer containing two α-chains and two β-chains with both α-chains linked to each other and to one β-chain, through disulfide bridges located at residues βC105 and αC72 for the α-β chain connections, and at residues αC15 and αC15 for the α-α chain connection (Figure 1A). In addition to those interchain disulfide bonds, each β-chain contains two intrachain disulfide bonds at residues C148-C179 and C190-C219, and each α-chain contains one intrachain disulfide bond at residues C34-C68. Four glycans of Hp 1-1 are attached to each β-chain (N23, N46, N50, and N80).

VISTA consists of 180 residues with 6 N-glycosites at residues N17, N59, N76, N96, N103 and N158. VISTA also contains 3 intra disulfide bonds located at residues C10-C146, C22-C114 and C51-C113.

Transferrin contains 679 residues with 2 N-glycosites at residues N413 and N614 and 19 disulfide bonds located throughout its structure.

To achieve maximal sequence coverage, HDX-MS analysis of each protein was performed with different concentrations of TCEP and urea in the quench buffer to facilitate denaturation and reduction of disulfide bonds.

For Hp 1-1, without online deglycosylation, a sequence coverage of 91.6% was achieved for the α-chain and a sequence coverage of 84.5% was achieved for the β-chain (Table 1, Figures S6). For the β-chain, 66 peptides could be identified, however, without coverage of any of the four N-linked glycosite regions (N23, N46, N50, and N80). At the N80 N-linked glycan site, two peptides were identified corresponding to the non-glycosylated form.

**Table 1:** Overview of glycoproteins subjected to HDX-MS analysis with the PNGase Rc column.

| Glycoprotein | Sequence coverage (%) without PNGase Rc column | No. of peptides (without PNGase Rc column) | Sequence coverage (%) with PNGase Rc column | No. of peptides (with PNGase Rc column) | No. of N-linked glycosites | No. of deglycosylated N-linked glycosites | Quench conditions (after 1:1 dilution) |
| --- | --- | --- | --- | --- | --- | --- | --- |
| c-Met | 91 | 721 | 98 | 755 | 6 | 6 | 125mM TCEP and 1.7M Urea pH 3 |
| Haptoglobin 1-1 (α Chain) | 92 | 9 | 100 | 100 | 0 | 0 | 125mM TCEP and 2M Urea pH2.3 |
| Haptoglobin 1-1 (β Chain) | 84.5 | 66 | 98 | 81 | 4 | 4 | 125mM TCEP and 2M Urea pH2.3 |
| Transferrin | 65 | 87 | 71 | 92 | 2 | 2 | 100mM TCEP and 2M Urea, pH 2 |
| VISTA | 73 | 50 | 97 | 110 | 6 | 6 | 200 mM TCEP and 2M Urea pH 2.6 |

In comparison, during HDX-MS with the PNGase Rc column, sequence coverage for the α-chain was 100%, and the sequence coverage for the β-chain was 98.8% (Figure 3 and Table 1). For the β-chain, 86 peptides with a total redundancy of 5.33 were found. The most significant result was the entire coverage of all four glycosites with substantial redundancy at glycosites N46, N50, and N80 (Figure 3).

**Figure 3:**
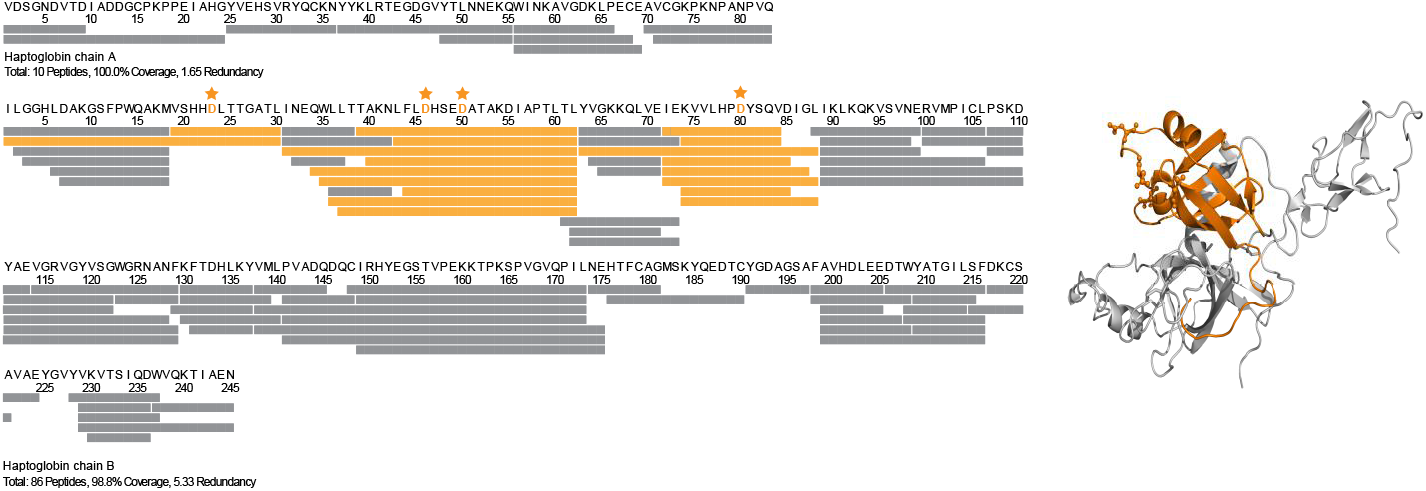
Sequence coverage map of Hp 1-1 obtained during HDX-MS with online deglycosylation by PNGase Rc. N-linked glycosites are highlighted in orange and marked by stars. Deglycosylated peptides are coloured in orange.

The HDX of Hp 1-1 was largely in agreement with its AlphaFold predicted structure. The regions around N46, N50, and N80 demonstrated rapid deuterium uptake, in contrast to the region around N23, which displayed slow HDX across the 10-hour time course (Figure S7 and S8). This is also consistent with where the N-glycan sites are situated inside the β-chain structure with N23 being situated within a β-strand. The solid hydrogen bonding of this region creates a more protected and stable area, which slows HDX. Conversely, all other N-glycosites are located in loop regions, which are characterized by being flexible explaining their faster HDX (Figure S7 and S8).

Similarly, HDX-MS analysis with the PNGase Rc column of VISTA and transferrin also showed a complete deglycosylation of all N-glycosites in each protein (Figure 4, Figure S9 and Table 1). The HDX of both VISTA and transferrin was in good overall agreement with their crystal structures with N-glycosites located in flexible loops exchanging more rapidly than those located in the more stable β-strands (Figures S10 and S12). Overall, the HDX-MS results on these three disulfide-bonded glycoproteins showed that incorporation of the PNGase Rc column in each HDX-MS experiment resulted in increased sequence coverage, redundancy and with complete coverage of all N-glycosites in each of the studied glycoproteins.

**Figure 4:**
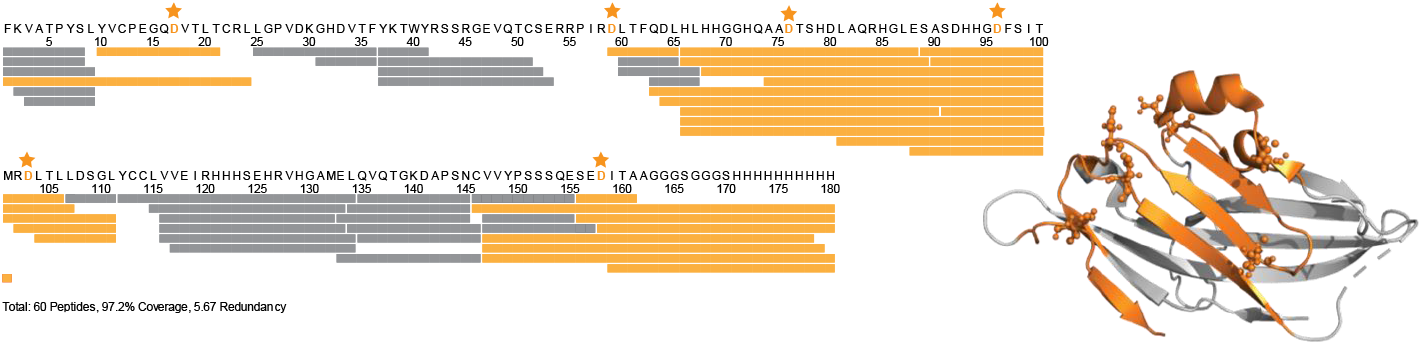
Sequence coverage map of VISTA during obtained during HDX-MS with online deglycosylation by PNGase Rc. N-linked glycosites are highlighted in orange and marked by stars. Deglycosylated peptides are coloured in orange.

The loss of deuterium label during HDX-MS (back-exchange) was monitored during analysis of Hp 1-1 with and without online deglycosylation (Figure S5). Based on the measurement of the deuterium contents of all detected peptides of Hp 1-1, we found that incorporation of the PNGase Rc column into the HDX-MS setup did not affect back-exchange (<2%).

### Epitope mapping by HDX-MS with online deglycosylation

We next applied HDX-MS with the integrated PNGase Rc column to map the binding epitope of a mAb on the complex glycoprotein c-Met. c-Met, (Mesenchymal Epithelial Transition factor), also named Hepatocyte Growth Factor Receptor (HGFR) is generated by the c-Met proto-oncogene and is regarded as a tumor assisted antigen and is an important signaling receptor for the tumor proliferating protein HGF (hepatocyte growth factor)^22^. Inhibition of c-Met has been indicated as a promising strategy to treat certain cancer forms^23^. The epitope mapping experiment was performed on the extracellular domain of c-Met (525 amino acids) that comprises an α- and a β chain, with the α chain containing 4 N-glycans and 4 intrachain disulfide bonds and the β chain containing 2 N-glycans and 1 intrachain disulfide bond. The two chains are connected by a disulfide bond between the C274 (chain α) and C36 (chain β).

The PNGase Rc-enabled HDX-MS epitope mapping experiment was done using a quench buffer containing TCEP and urea as this had previously been shown to yield the maximum sequence coverage of c-Met^16^. The HDX-MS experiment allowed identification and extraction of HDX information from 124 peptides covering 96.8% of c-Met. Importantly, this included peptides covering all 6 N-glycosites. Overall, the HDX of c-Met was in good agreement with the available crystal structure, with slow HDX in regions of stable structure and faster exchange in loops (Figure S11). For the six glycosylated regions, slow HDX was observed in regions 41-50, 102-117 and 196-218 indicating the presence of significant secondary structure, with fast HDX regions 83-95, 96-100 and 147-159 displaying HDX corresponding to highly solvent exposed loops. Upon binding the mAb, significant reductions in HDX was observed in several peptides spanning the region of the N202-glycosite. None of the other regions of c-Met, including the 5 other N-glycosylated regions showed significant changes in HDX (Figure 5B). Region 196-218 is thus clearly linked to binding of the mAb and it is very likely that the epitope is contained in this region. Notably, this region was previously inaccessible for HDX-MS analysis without the incorporation of the PNGase Rc column to allow efficient deglycosylation. A visual representation of the epitope when overlaid onto the crystal structure of c-Met (Figure 5C) shows that the region is highly solvent exposed and several residues are well-positioned for direct interaction with the mAb.

**Figure 5:**
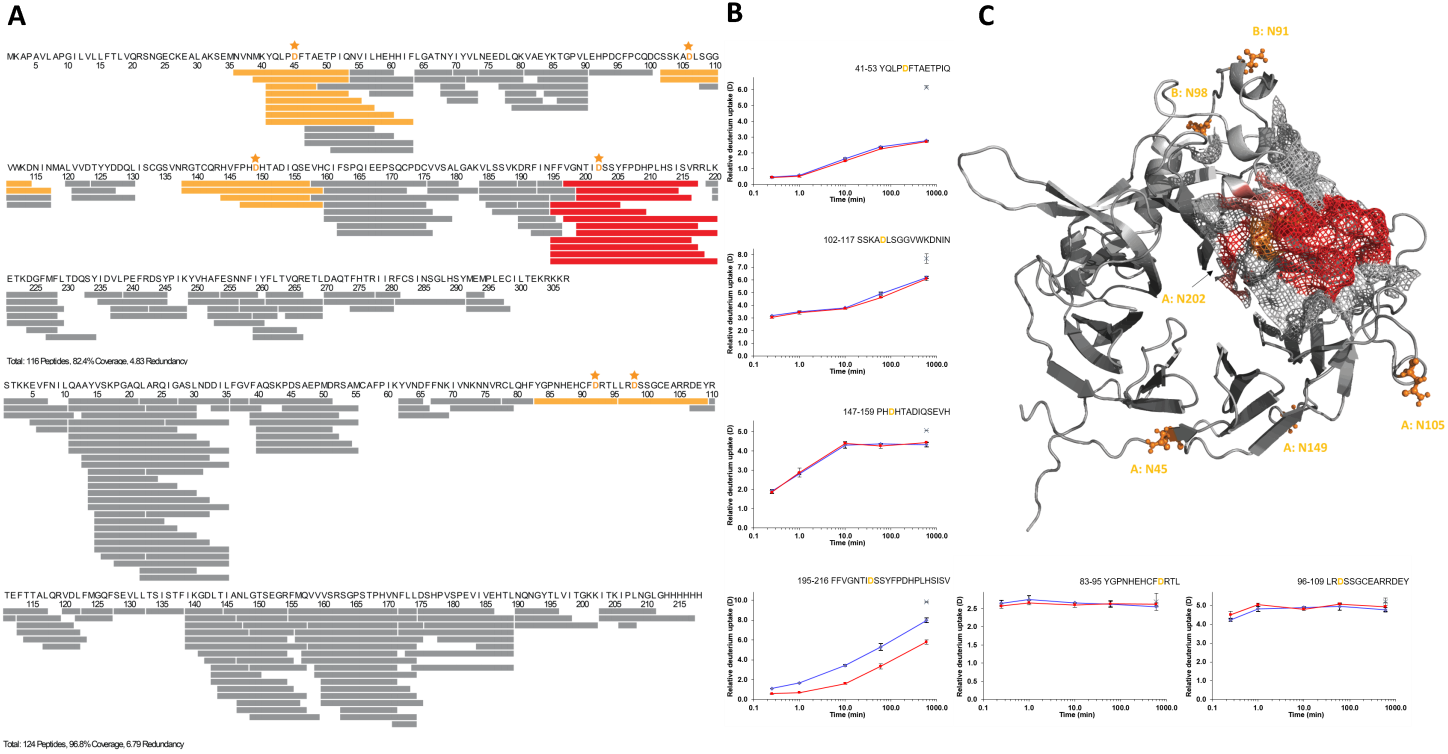
HDX-MS epitope mapping of c-Met with online deglycosylation. A) Sequence coverage map of c-Met, with deglycosylated peptides that contain an N-linked glycosite (denoted by orange star) highlighted in orange. Peptides highlighted in red showed decreased HDX upon mAb binding. B) HDX uptake plots of all deglycosylated peptides spanning an N-linked glycosite. c-Met with mAb uptake is highlighted in red, c-Met without mAb is highlighted in blue and the maximally-labelled control sample is shown as a cross. Error bars represent the standard deviation from three technical replicate measurements. C) Overlay of the HDX-MS data onto the crystal structure of c-Met (PDB: <u>1SHY</u>) with the epitope region displaying reduced HDX upon mAb binding (res. F-195 -V-216) shown in red and N-glycosites highlighted as orange sticks.

### Reusability and activity assessment of the PNGase Rc column

Reusability and activity of the PNGase Rc column was measured through the developed LC-MS assay using tryptic glycopeptides from the IgG1 Trastuzumab as substrate^16^. To investigate the activity of the PNGase Rc column at different temperatures, the enzymatic activity was measured at 0°C and room temperature. A reduced activity of 33% was observed at 0°C (Figure 6B). However, a high degree of deglycosylation could still be obtained at 0°C, which is suitable for an HDX-MS experiment if required. Notably, the HDX-MS experiments for Hp 1-1, transferrin, c-Met were all done with the column placed outside the cooling box at room temperature whereas the HDX-MS experiment of VISTA was performed with online deglycosylation at 0°C.

**Figure 6:**
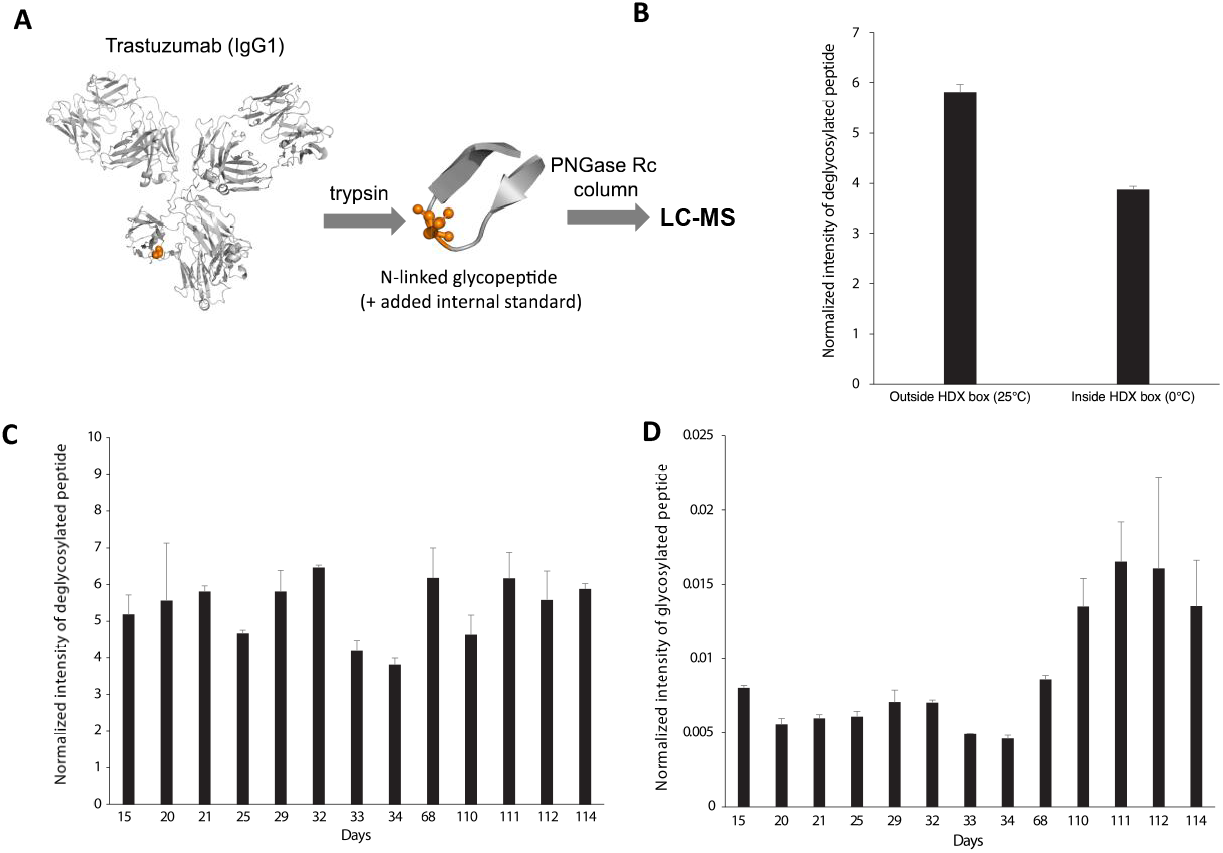
Monitoring the reusability and temperature dependence of the PNGase Rc column. A) LC-MS assay used to monitor the activity of the PNGase Rc column. Intensities of the IgG1 N-linked glycopeptide EEQYNSTYR were normalized to intensities of the internal standard Angiotensin II. B) Activity of the PNGase Rc column when placed inside (*0°C)* and outside the HDX box (*20°C*) at a flow rate of 200 μl/min. C) Reusability of the PNGase Rc column over 114 days (monitored by LC-MS assay). During this period, the PNGase Rc column was used regularly in various HDX-MS experiments exposing it to TCEP and urea (see Table 1). The column was used and equilibrated from day 0 to day 15.

Next, to investigate the reusability of the PNGase Rc column, the intensity of the glycosylated and deglycosylated form of the glycopeptide was monitored by LC-MS at regular intervals over a time period of 114 days. During this time period, the column was regularly used for other HDX-MS experiments exposing it to TCEP and urea. Overall, the performance and stability of the column was very good. The fluctuations in intensity of the glycosylated peptide around day 34 to 68 could be caused by changes in substrate batches on day 68 and 110. Nevertheless, from both the intensities of the deglycosylated and the glycosylated peptide, it is clear that even after 114 days of regular usage of the PNGase Rc column, the overall deglycosylation efficiency remains high and largely unchanged (Figure 6C-D). Importantly, this included regular exposure to urea and TCEP over the whole time period. The tolerance of the PNGase Rc column to chaotropic - and reducing agents is of high importance, since glycoproteins are often extracellular, and thus very often highly disulfide bonded^24^ and thus require both chaotropic and reducing agents in the quench buffer during HDX-MS^5,16^

Furthermore, the same LC-MS assay was used to investigate the deglycosylation efficiency of the PNGase Rc column while using quench buffers with high concentrations of TCEP and urea (2.5M and 3M urea and 200mM and 400mM TCEP). The results show that exposure to such high concentrations of TCEP and urea results in some decrease in deglycosylation efficiency compared to the reference (no TCEP and urea) (Figure 7). Importantly, this decrease in deglycosylation efficiency was fully reversible. Furthermore, as shown in the prior section, the column was fully capable of comprehensive deglycosylation of the different glycoproteins during HDX-MS experiments using quench buffers containing substantial amounts of TCEP and urea (Table 1). Thus, the very good reusability and tolerance to urea and TCEP should make the developed PNGase Rc column of significant utility in improving the HDX-MS analysis of glycoproteins.

**Figure 7:**
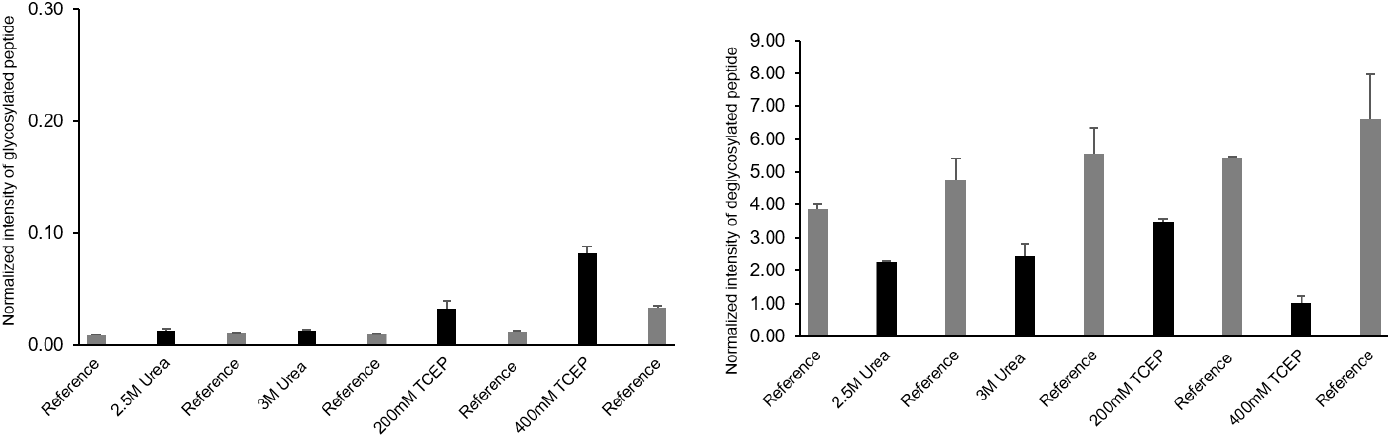
Deglycosylation efficiency of the PNGase Rc column in the presence of different concentrations of TCEP and urea. The intensity of the glycopeptide EEQN/DSTYR was monitored in its glycosylated and deglycosylated form and was normalized to an internal standard. The reference condition represents deglycosylation of the glycopeptide under identical conditions but without the addition of TCEP and urea.

We note that at the same time of this work^25^, a paper describing the successful development of a PNGase Dj column for use in HDX-MS has been reported^26^ This work extends our earlier work on immobilization of PNGase Dj in a microchip to a column format. It thus appears that both PNGase Rc and Dj can be immobilized on-column to advance analysis of glycoproteins by HDX-MS.

## CONCLUSIONS

Here we show that the developed PNGase Rc column enables highly efficient online removal of N-linked glycans during HDX-MS of glycoproteins. Implementation of the PNGase Rc column to the conventional HDX-MS setup in each lab was straightforward, placing it immediately downstream from the pepsin column to allow for efficient online digestion and deglycosylation. HDX-MS experiments were performed on four complex glycoproteins that required harsh quench conditions (up to 2M urea and 200mM TCEP) including Hp 1-1 (4 N-linked glycans, 7 S-S bonds), VISTA, 6 N-linked glycans, 3 S-S bonds), transferrin (2 N-linked glycans, 4 S-S bonds) and c-Met (6 N-linked glycans, 5 S-S bonds). In all cases, significantly increased sequence coverage was obtained through use of the PNGase Rc column, enabling measurement of the HDX of all glycosylated regions that were previously inaccessible without the column. We also show how the PNGase Rc column enables epitope mapping of an epitope on c-Met, a challenging glycoprotein drug target. Control experiments showed that use of the PNGase Rc column during HDX-MS resulted in negligible increases (<2%) in the loss of deuterium label (back-exchange).

The column retains high activity in the presence of common quench-buffer additives like TCEP and urea. A moderate reduction in activity was observed at high concentrations of TCEP (200-400mM TCEP) and urea (2.5-3M), but importantly this reduction was fully reversible. Column performance was consistent across repeated injections containing TCEP and Urea and stable for >100 days.

We envision that the straight-forward incorporation of the PNGase Rc column into conventional HDX-MS setups could greatly enable HDX-MS analysis of the conformation and ligand interactions of many challenging glycosylated proteins that have so far presented a challenge to HDX-MS, including numerous attractive and unexplored drug targets. Additionally, the column could also be used in other bottom-up proteomics workflows that can benefit from fast online deglycosylation of peptides or proteins prior to LC separation and MS. The developed PNGase Rc column is commercially available (AffiPro, https://AffiPro.cz/).

## ASSOCIATED CONTENT

### Supporting Information

Supporting Information is available and includes: immobilization protocol, immobilization efficiency test protocol, test of nonspecific adsorption to POROS 20 AL beads, PNGase Rc AlphaFold structure and active site prediction, activity of immobilized PNGase Rc, coverage maps of both glycosylated and deglycosylated Hp 1-1, HDX uptake plots of Hp 1-1, VISTA and transferrin, maximum-labeled control data for Hp 1-1 and HDX uptake plots for the c-Met epitope experiment.

## AUTHOR INFORMATION

### Author Contributions

KDR, AZ, MK designed the study. PK and DD produced PNGase Rc. MG, PP and PN performed immobilization with efficiency and enzyme activity tests. PN performed SDS-PAGE analysis of deglycosylation of Hp 1-1. Luisa S performed HDX-MS of Hp 1-1. MG performed HDX-MS of VISTA. Luke S performed HDX-MS of transferrin. TL and CW performed HDX-MS epitope mapping of c-Met. CW, UR, JB, AZ, KDR supervised the study. The draft of the manuscript was written by T.L and KDR – and revised through contributions of all authors. All authors have given approval to the final version of the manuscript.

## ACKNOWLEDGMENT

This work was supported by the European Research Councils (ERC Consolidator Grant no. 101003052, to K.D.R.) and the Roche Biologics Technology Evaluation & Development program. HDX-MS analysis of VISTA was performed using an RSLC U3000 HPLC system and an Orbitrap Eclipse Tribrid Mass Spectrometer financed by the European Regional Development Fund (ERDF) and the State Ministry of Baden-Wuerttemberg for Economic Affairs, Labor and Tourism (#3-4332.62-NMI/69). We acknowledge the Structural mass spectrometry facility of CIISB, Instruct-CZ Centre, supported by MEYS CR (LM2023042) and European Regional Development Fund-Project "UP CIISB“ (No. CZ.02.1.01/0.0/0.0/18_046/0015974).

