## Supplementary figures for "Development of a PNGase Rc column for online deglycosylation of complex glycoproteins during HDX-MS"

This PDF file includes:

Materials and methods
Figs.
Tables

Supporting Information

### 1 Section 1

PNGase Rc expression and purification. Expression of PNGase Rc was done as described earlier[18]. A 50 ml starter culture was added to a 1L flask containing LB medium and, the antibiotic kanamycin and incubated at 37°C with shaking at 160 RMP. Samples of 1mL were taken at time points 0, 30, 60, 90, and 120 min, and the optical density was measured at 600nm with a UV-spectrophotometer. Induction was done when the OD600 had reached 0.6 to 0.8, the protein expression was induced by adding 1mL of 1mM IPTG and incubated 1 hour. The temperature was lowered to 18°C and the expression culture was then left to incubate overnight before harvesting the cells by centrifugation at 6000 rpm for 20 min at 4 °C.

The supernatant was discarded and the cell pellets were resuspended in lysis buffer pH 7.5 (100mM TRIS, 1% Triton X-100 (v/v) and 1mM phenyl-methylsulfonyl fluoride (PMSF)) and sonicated for 20 min (40 on/off cycles with 20 µm amplitude for 15 s at 4°C). The lysate was then centrifuged at 10000 rpm for 20 min at 4 °C. Pellets were discarded and the supernatant was loaded onto a pre-washed and -equilibrated Ni-NTA column. 50mM imidazole was used to remove unspecifically bound species and protein was eluted with 500mM imidazole. Eluted fractions were pooled and purified further on a size-exclusion column (Sephadex G-25) using running buffer of 20mM sodium phosphate dibasic, 500mM NaCl. Purified PNGase Rc (>90% as assessed by SDS-PAGE) was stored in aliquots at -80°C.

Larger scale production of PNGase Rc was performed according to a similar protocol described earlier[19].

Tryptic digestion of Trastuzumab.

TZ (1500pmol) was denatured in 6 M guanidinium hydrochloride (GdnHCl) in 50mM NH 4HCO3 pH 8 at 60 °C for 30 min. Disulfide bond reduction was carried out by the addition of 20µL of 45mM dithiothreitol (DTT) at 60 °C for 2 h, 20µL of 45mM iodoacetamide was added followed by incubation at room temperature (RT) for 30 min. The mixture was diluted by the addition of 400µL of 50mM $NH_4HCO_3$ pH 8, and trypsin was added at a 1:20 w/w ratio with TZ. The digestion took place overnight at 37 °C. Each sample aliquot was prepared by adding 10µl of the tryptic digest along with 1µl of 10pmol Angiotensin II (ATII) to 89µl 0.23% formic acid (pH2.3). Each sample injection thus contained 50pmol of TZ digest glycopeptides and 10pmol of ATII as the internal standard[16].

### 2 Immobilization of PNGase Rc to POROS$^{TM}$ 20 AL beads

#### 2.1 Prediction of immobilization sites

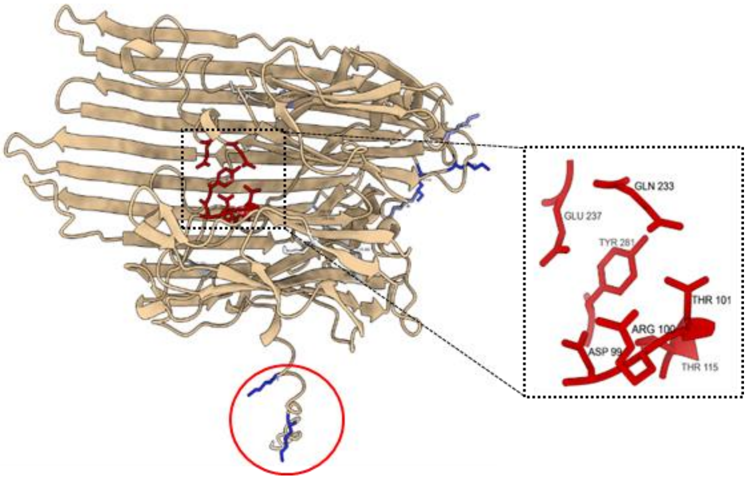

*Figure S1: PNGase Rc AlphaFold structure. Residues in red belong to the predicted active site in PNGase Rc. Residues in blue are lysine amino acids. SASA of lysine is represented by the degree of the blue. The darker the blue, the higher is the residue exposed to the solvent. Lysines in the red circle are more likely immobilization sites.*

#### 2.2 Immobilization method description

Reagents and consumables

- 9.3 mg (3.6 mg/mL) PNGase Rc produced as described in reference 17.
- 93 mg POROS$^{TM}$ 20 AL Aldehyde activated Resin (Thermofisher, #1602906)
- Trastuzumab (kindly domated by MAB Discovery GmbH, (Neuried, Germany))
- $NaCNBH_3$ (5M in 1M NaOH, SigmaAldrich, #296945)
- Ethanolamine, sodium azide, citric acid, sodium hydroxide (SigmaAldrich)
- pepsin column (Enzymate BEH Pepsin Column, Waters, Eschborn, Germany)
- VISTA (Human B7-H5, Acro BioSystems)

For covalent PNGase Rc immobilization, first the 5 M NaCNBH3 commercial stock solution in 1 M NaOH (SigmaAldrich, #296945) was diluted 1:5 with 150 mM citrate/NaOH buffer pH 2.5 to give a final concentration of 1 M NaCNBH3 at pH 7.5. 150 µL of this NaCNBH3 solution were mixed with 4.7 mg PNGase Rc (1.3 mL, 3.6 mg/mL in PBS buffer) and 47 mg POROSTM 20 AL aldehyde activated resins (20 µm particle size with throughpores and diffusive pores for perfusion chromatography, Thermofisher, #1602906) to give an enzyme-to-bead ratio of 1:10. The bead slurry was incubated over night at room temperature in an overhead shaker. Afterwards, the slurry was centrifuged at 1000 x g for 1minute, the supernatant was decanted and 50 µL of supernatant were taken for determination of the coupling efficiency of PNGaseRc to the beads. The reaction was stopped and free aldehydes were blocked with 1.5 mL ethanolamine solution (1M, pH 7.4) by incubation for 30 minutes at RT in an over-head shaker. Beads were washed five times with 1 mL PBS and supernatant was discarded each time after centrifugation. Beads were stored as 50% slurry in PBS containing 0.02 % sodium azide at 4 °C.

#### 2.3 Immobilisation Efficiency test

Immobilization efficiency (ψ) was tested by assessment of PNGase Rc concentration in the supernatant after bead coupling $\psi(\%)=(conc._{Initial}-conc._{Supernatant})/conc._{Initial}*100$. The concentration was determined from a calibration curve using the UV280nm peak area of standard dilutions of PNGaseRc (see Figure S2) analyzed by the RP-C4-LC-UV method described in Gramlich et al., Anal. Chem. 2022, 94, 27, 9863–9871. Prior to analysis of the supernatant, NaCNBH3 was removed by ultrafiltration using Amicon filters (10k MWCO, MerckMillipore). In a separate experiment, it was shown that ultrafiltration leads to a loss of 13% PNGase Rc. This loss was taken into account for determination of the immobilization efficiency.

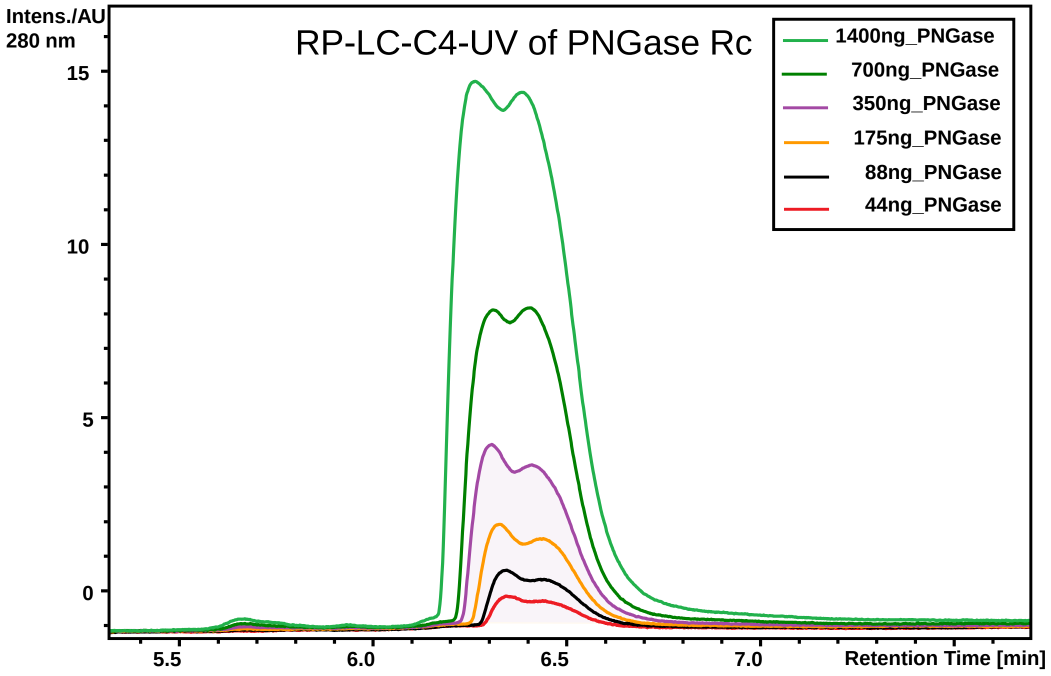

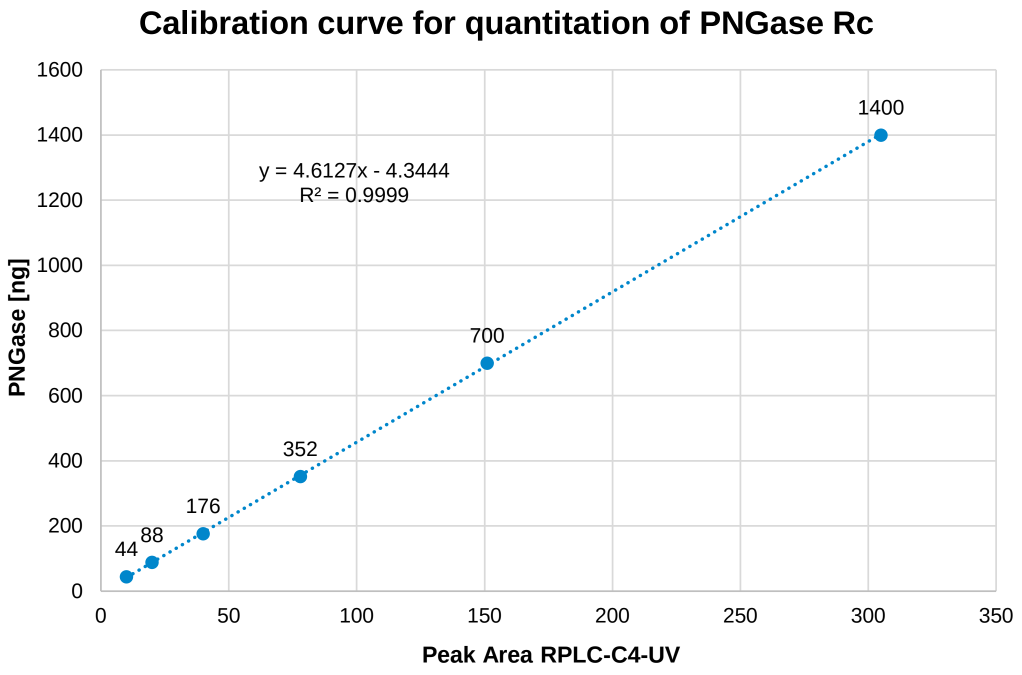

*Figure S2: RPLC-C4-UV used for determination of the degree of immobilization of PNGase Rc to POROS$^{TM}$ 20 AL beads.*

An enzyme-to-bead (E:B) ratio of 1:10 yielded ~ 39% immobilization efficiency, whereas an E:B ratio of 1:20 yielded 95% immobilization efficiency. Therefore, the E:B ratio of 1:20 results in quantitative coupling of the enzyme to the beads but most likely not the whole accessible surface is modified with the enzyme.

#### 2.4 Noncovalent adsorption of protein to POROS™ 20 AL beads

Nonspecific adsorption of protein to POROSTM 20 AL beads was tested using Trastuzumab and different incubation times with the beads (Figure S3). Very little loss of Tz was obtained after incubation with the beads for 30 minutes.

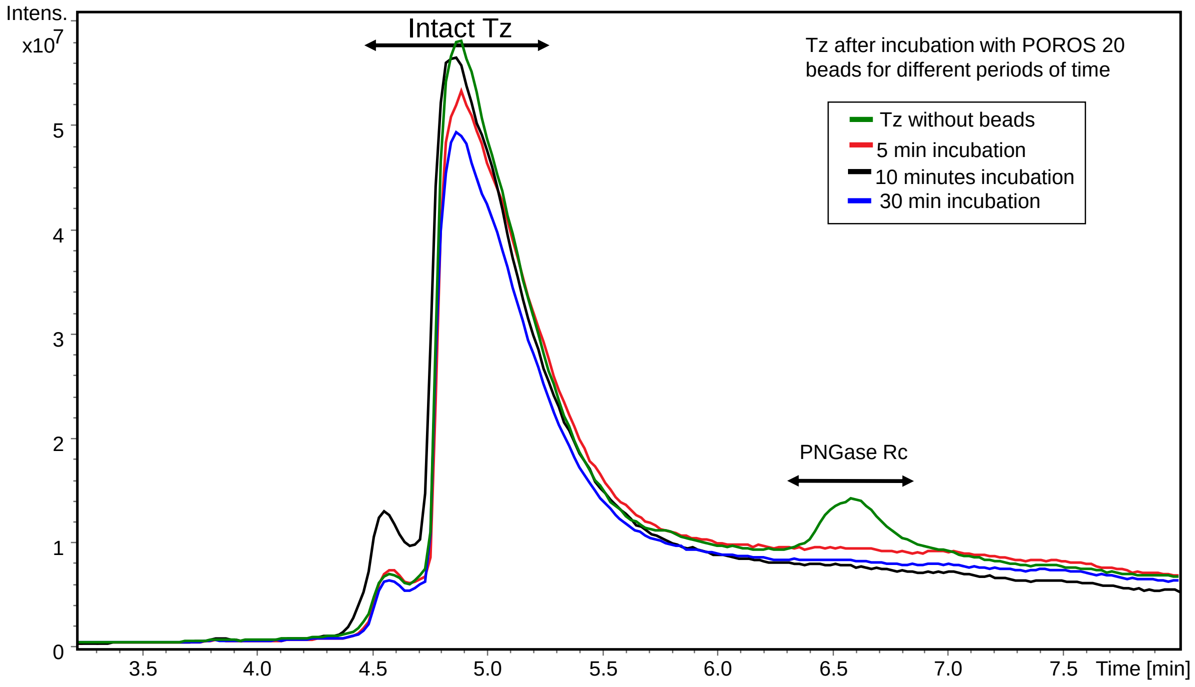

*Figure S3: Test of noncovalent adsorption of Tz to the POROS™ 20 AL beads by RPLC-C4-MS.*

#### 2.5 Test of PNGase Rc enzyme activity after immobilization

The enzyme activity test was performed as described in M.Gramlich et al[16]. For this, 10 µL bead slurry (50% in PBS) was centrifuged and supernatant was discarded. Deglycosylation was initiated by adding 45 µL of a solution of 5 µL Trastuzumab (Tz) (5 mg/mL) and 40 µL citrate/NaOH buffer (pH 2.5) to the beads. Samples were incubated for 5, 10 and 30 minutes by overhead mixing. Afterwards, they were centrifuged and 15 µL of the supernatant taken and quenched by heat for 5 minutes at 95°C. In the subsequent RP-C4-LC-MS analysis[16], the peak area obtained for trastuzumab (with and without deglycosylation) in the total ion chromatogram (TIC) after incubation with the modified beads for different periods of time was assessed (Figure S4). The enzyme activity assays showed 100 % deglycosylation for Tz for beads with an E:B ratio of 1:10 and 40 % deglycosylation for an E:B ratio of 1:20 (5 min. incubation, pH 2.5, RT). For comparison: PNGase Rc in solution showed 20 % deglycosylation of Tz under the same conditions.

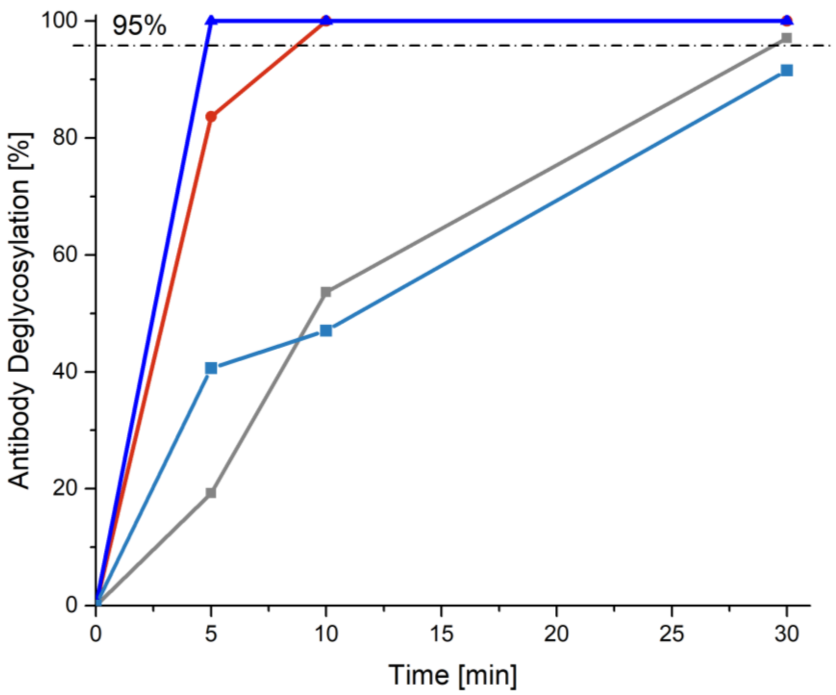

*Figure S4: Enzyme activity of immobilized PNGase Rc. In-solution enzyme activity (measured as described earlier[19]) highlighted in grey, bead batch 01: 1:10 E:B with solid $NaCNBH_3$ highlighted in red, bead batch 02: 1:10 E:B with stock sol. $NaCNBH_3$ highlighted in dark blue and bead batch 03: 1:20 E:B with stock sol. $NaCNBH_3$ highlighted in light blue.*

### 3 Assessment of HDX back exchange with and without online PNGase Rc column

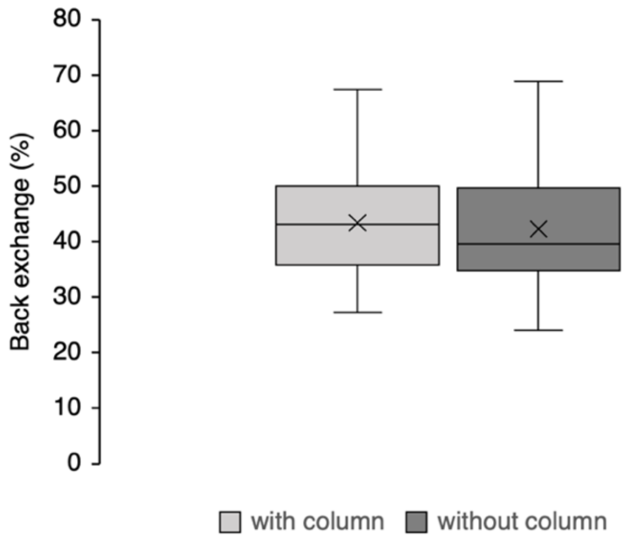

*Figure S5: Comparison of back exchange values of all identified peptides from Hp 1-1 during HDX-MS analysis with and without the the PNGase Rc column.*

### 4 Haptoglobin 1-1

#### 4.1 Sequence coverage without online PNGase Rc deglycosylation

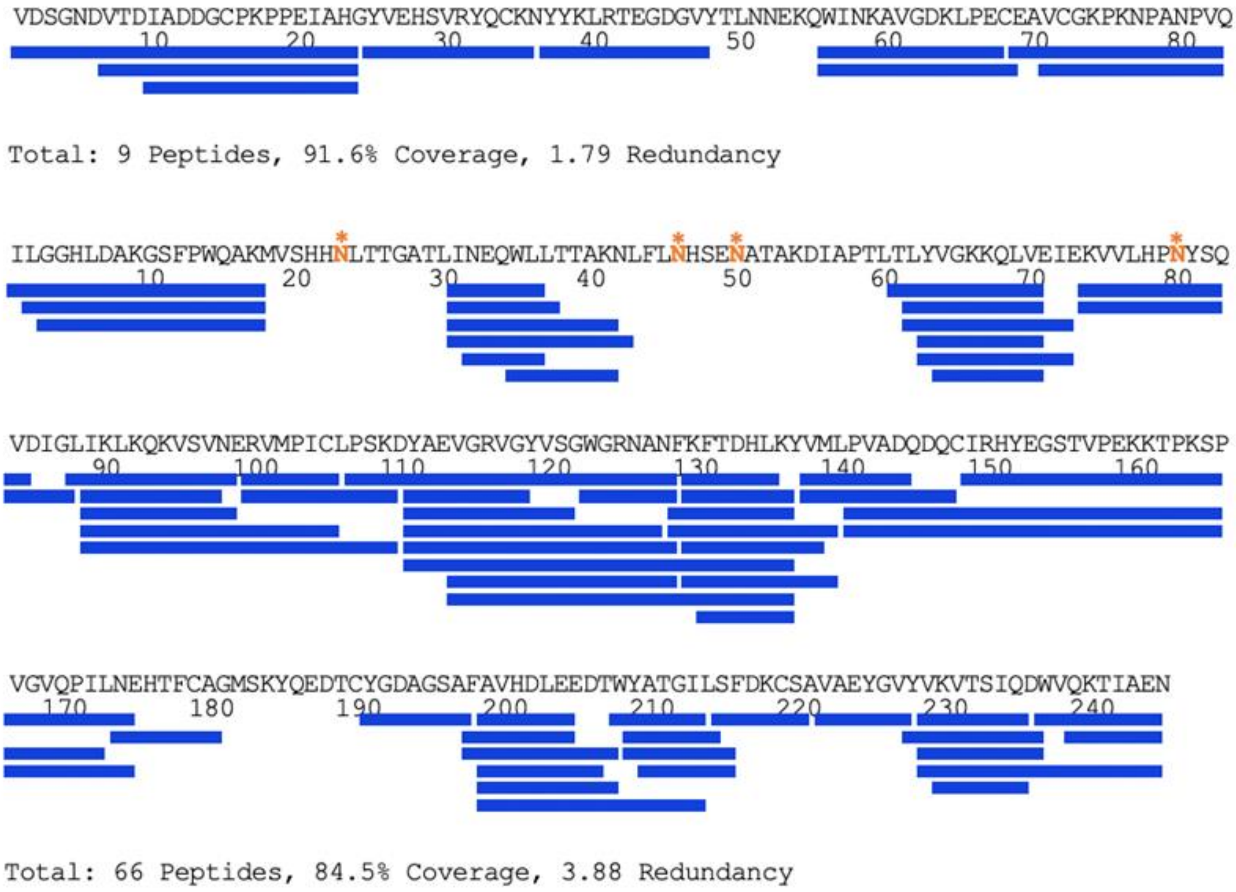

*Figure S6: Coverage map of glycosylated Hp 1-1 α-chain (top) and Hp 1-1 β-chain (bottom). Identified peptides are shown as blue bars and glycosylated Asn residues are highlighted in orange and marked with an orange star.*

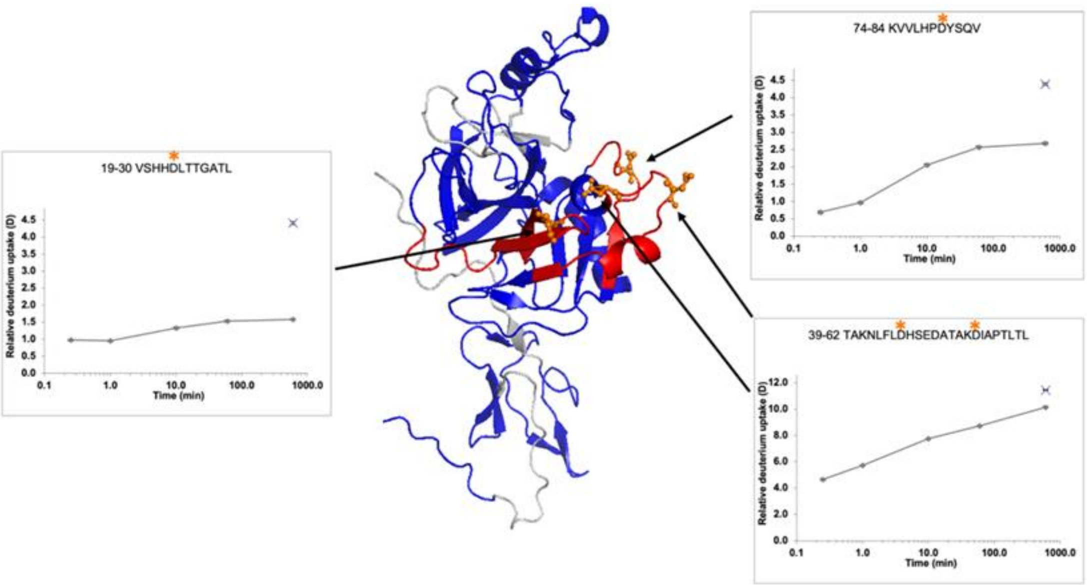

*Figure S7: Structure of the alpha and beta-chain of Haptoglobin 1-1. Covered regions are colored in blue, uncovered regions are shown in grey, and glycosylated residues are shown in orange. Uptake plots from a peptide of each N-linked glycan region are shown and are coloured red in the crystal structure. Maximally labelled control samples are shown in the uptake plots.*

#### 4.2 HDX uptake plots of Hp 1-1

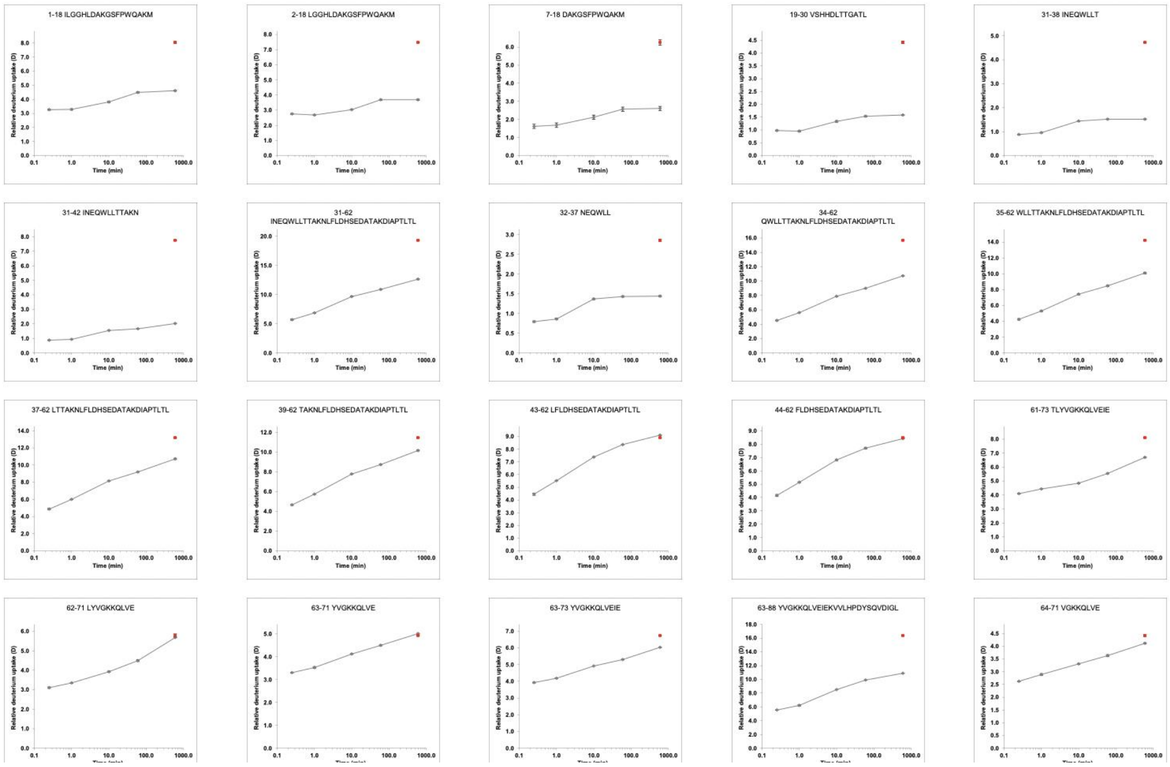

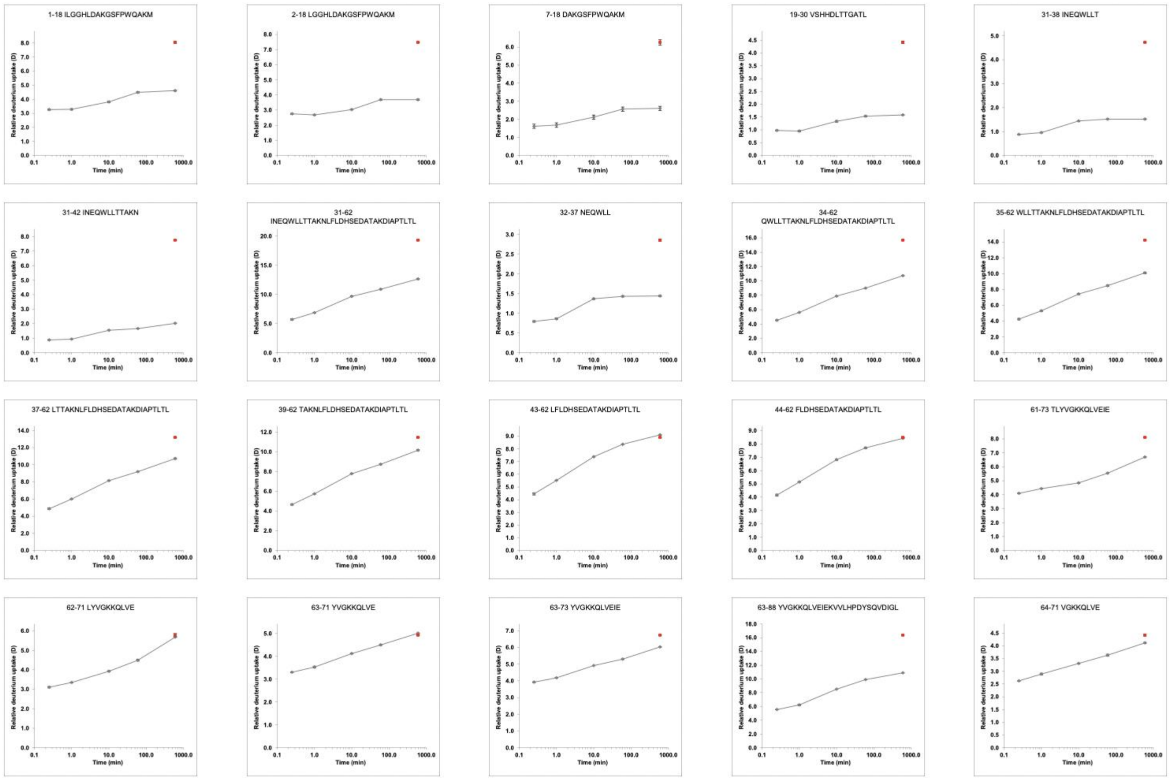

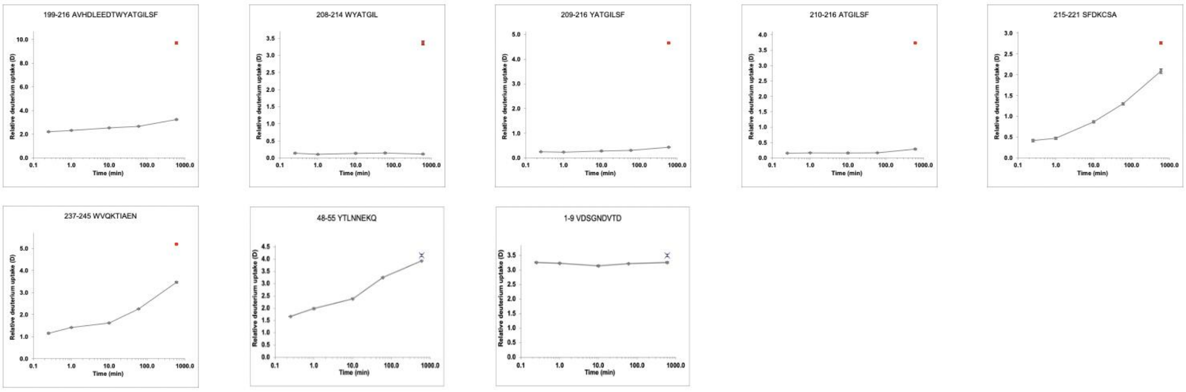

*Figure S8: HDX uptake plots of peptides of Hp 1-1. Error bars denote standard deviations from triplicate technical replicates. The deuterium content of the maximally-labeled sample is denoted by a red dot.*

### 5 Transferrin

#### 5.1 Sequence coverage with online PNGase Rc deglycosylation

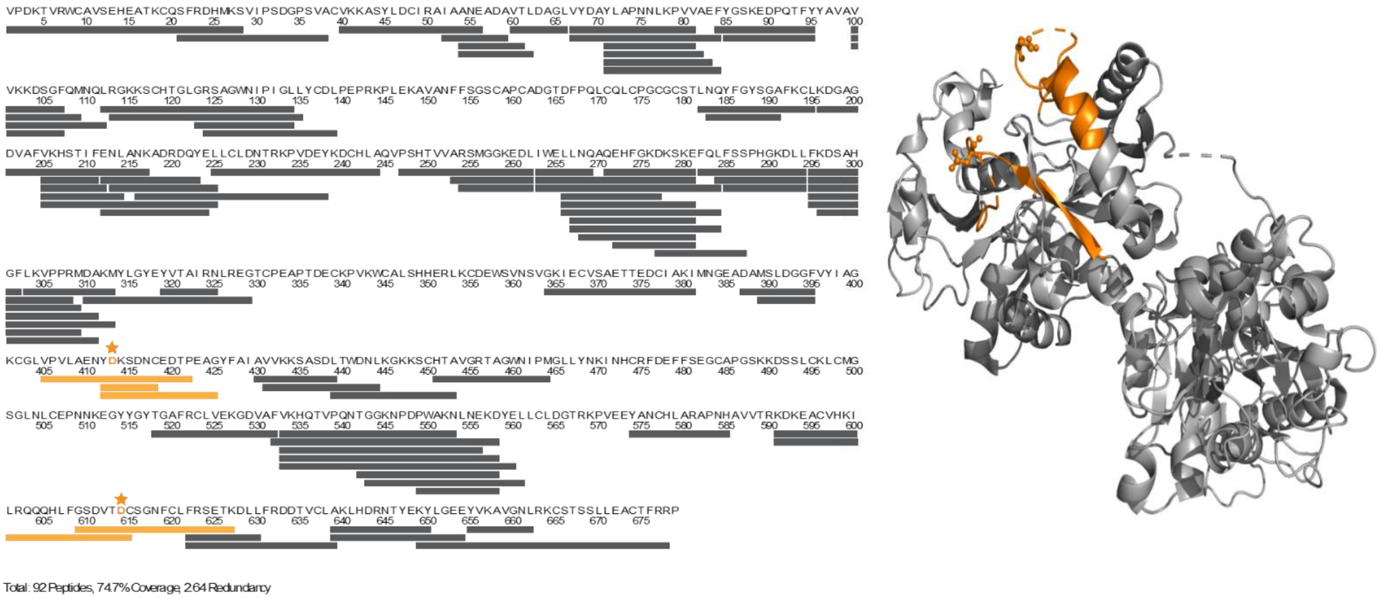

*Figure S9: Coverage map of Transferrin during HDX-MS with online deglycosylation. Deglycosylated peptides are highlighted in orange*

#### 5.2 HDX uptake plots of Transferrin

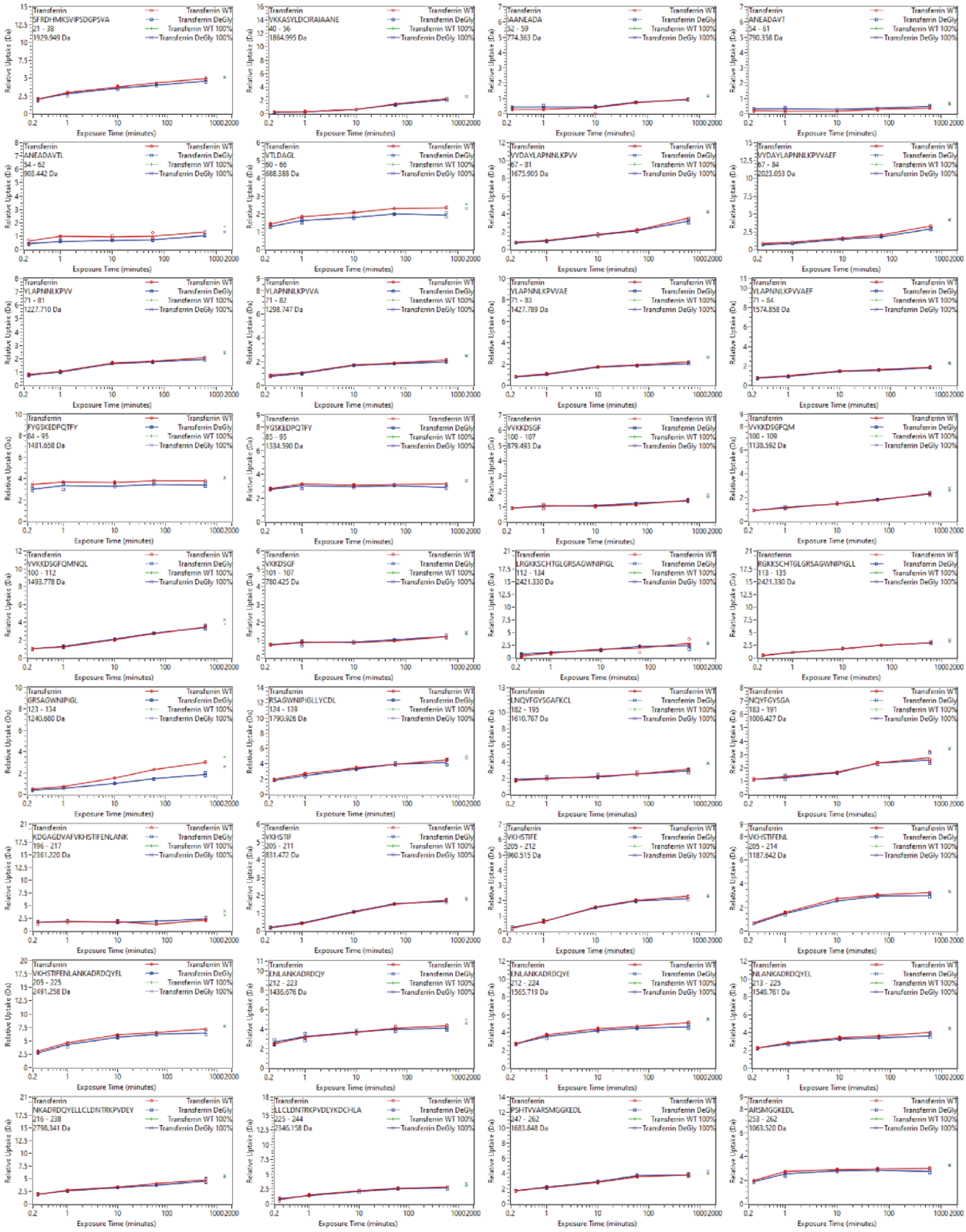

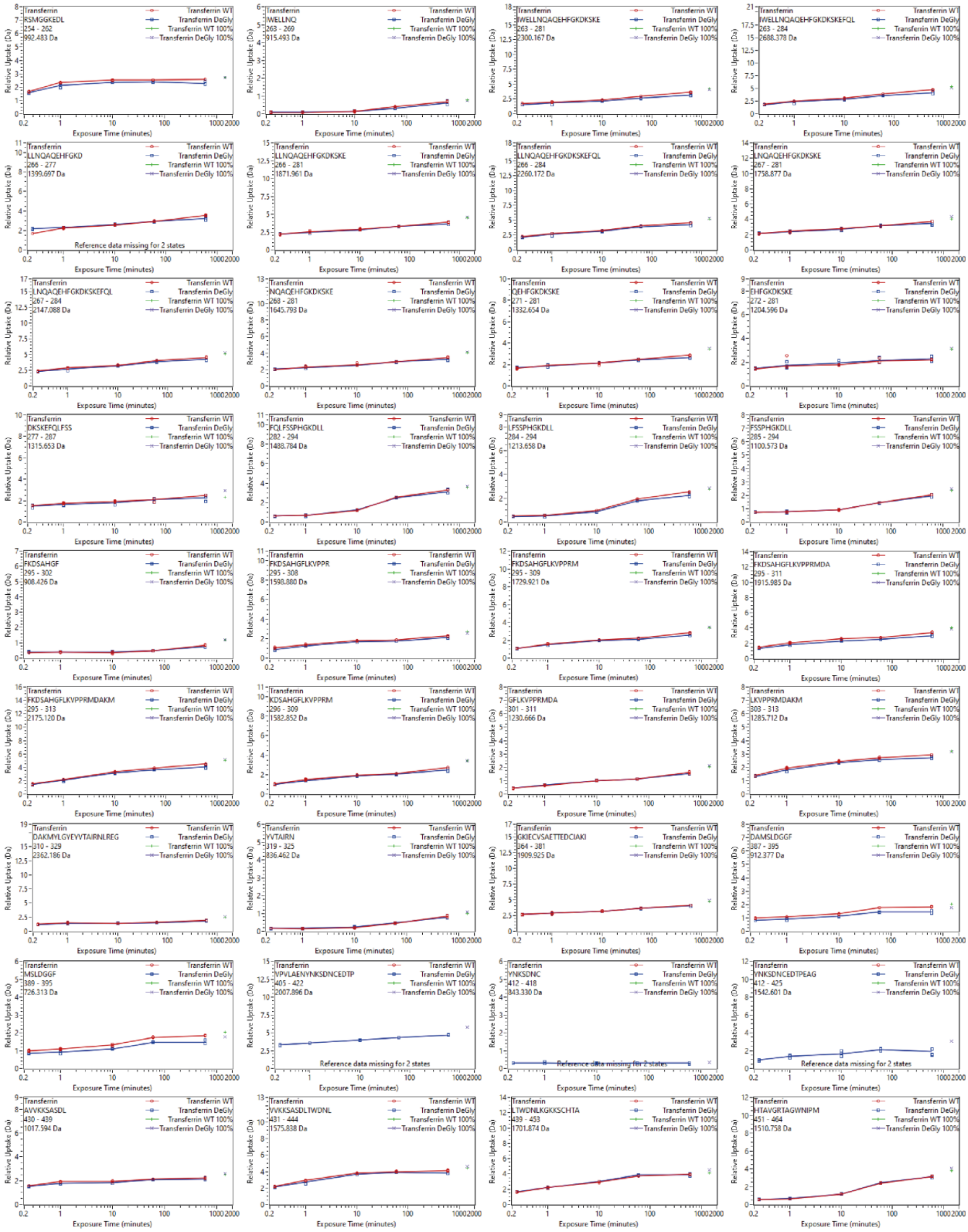
Transferrin
Transferrin WT
Transferrin DeGly
Transferrin WT 100%
Transferrin DeGly 100%
Relative Uptake (Da)
Exposure Time (minutes)
RSMGGKEDL 254 - 262
IWELLNQ 263 - 269
IWELLNQAQEHFGKDKSKE 263 - 281
IWELLNQAQEHFGKDKSKEFQL 263 - 284
LLNQAQEHFGKD 266 - 277
LLNQAQEHFGKDKSKE 266 - 281
LLNQAQEHFGKDKSKEFQL 266 - 284
LNQAQEHFGKDKSKE 267 - 281
LNQAQEHFGKDKSKEFQL 267 - 284
NQAQEHFGKDKSKE 268 - 281
QEHFGKDKSKE 271 - 281
EHFGKDKSKE 272 - 281
DKSKEFQLFSS 277 - 287
FQLFSSPHGKDLL 282 - 294
LFSSPHGKDLL 284 - 294
FSSPHGKDLL 285 - 294
FKDSAHGF 295 - 302
FKDSAHGFLKVPPR 295 - 308
FKDSAHGFLKVPPRM 295 - 309
FKDSAHGFLKVPPRMDA 295 - 311
FKDSAHGFLKVPPRMDAKM 295 - 313
KDSAHGFLKVPPRM 296 - 309
GFLKVPPRMDA 301 - 311
LKVPPRMDAKM 303 - 313
DAKMYLGYEYVTAIRNLREG 310 - 329
YVTAIRN 319 - 325
GKIECVSAETTEDCIAKI 364 - 381
DAMSLDGGF 387 - 395
MSLDGGF 389 - 395
VPVLAENYNKSDNCEDTP 405 - 422
YNKSDNC 412 - 418
YNKSDNCEDTPEAG 412 - 425
AVVKKSASDL 430 - 439
VVKKSASDLTWDNL 431 - 444
LTWDNLKGKKSCHTA 439 - 453
HTAVGRTAGWNIPM 451 - 464
Reference data missing for 2 states

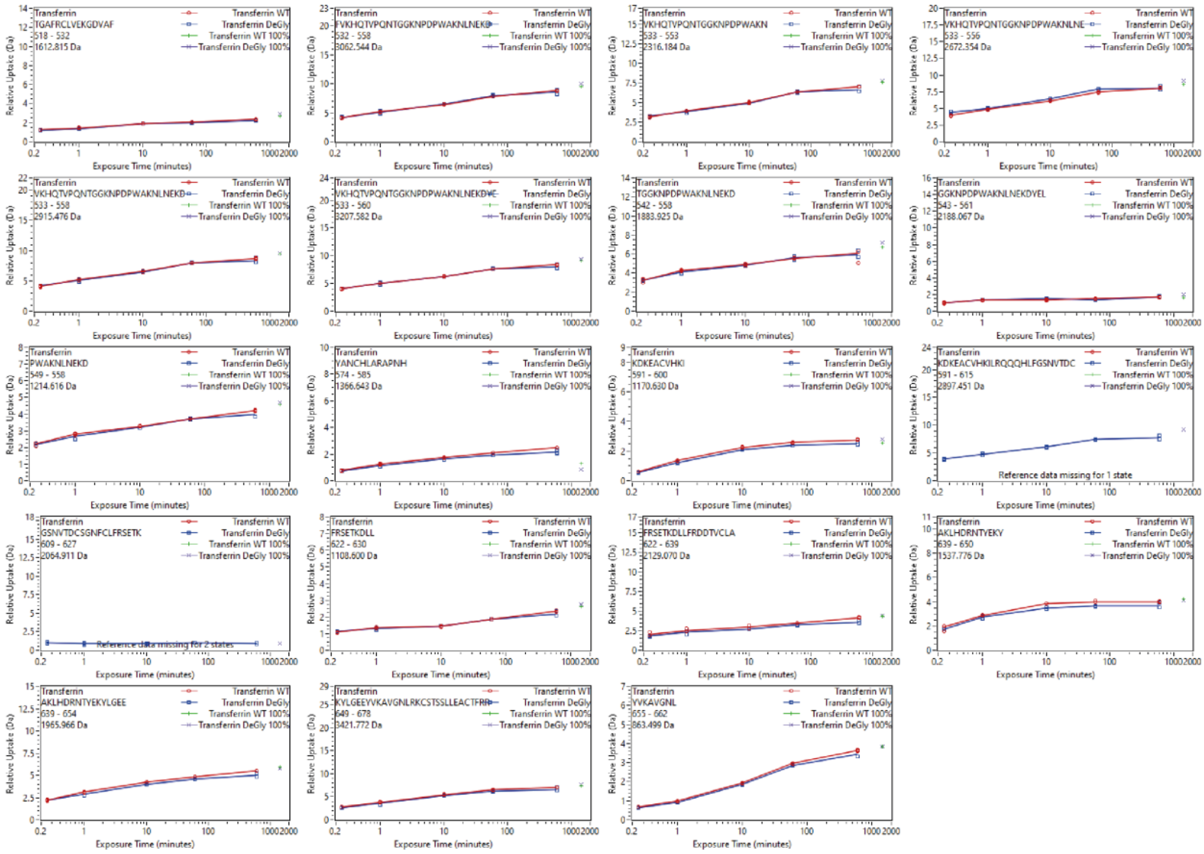

*Figure S10: HDX uptake plots of peptic peptides of Transferrin. The uptake of Transferrin WT is highlighted red, Transferrin post deglycosylation is highlighted blue and maximally-labelled control samples are cross marked green for WT and Purple for deglycosylated Transferrin. Error bars represent the standard deviation from three technical replicate measurements. Error bars denote standard deviations from triplicate technical replicates.*

### 6 HDX uptake plots of c-Met

#### C-Met chain α

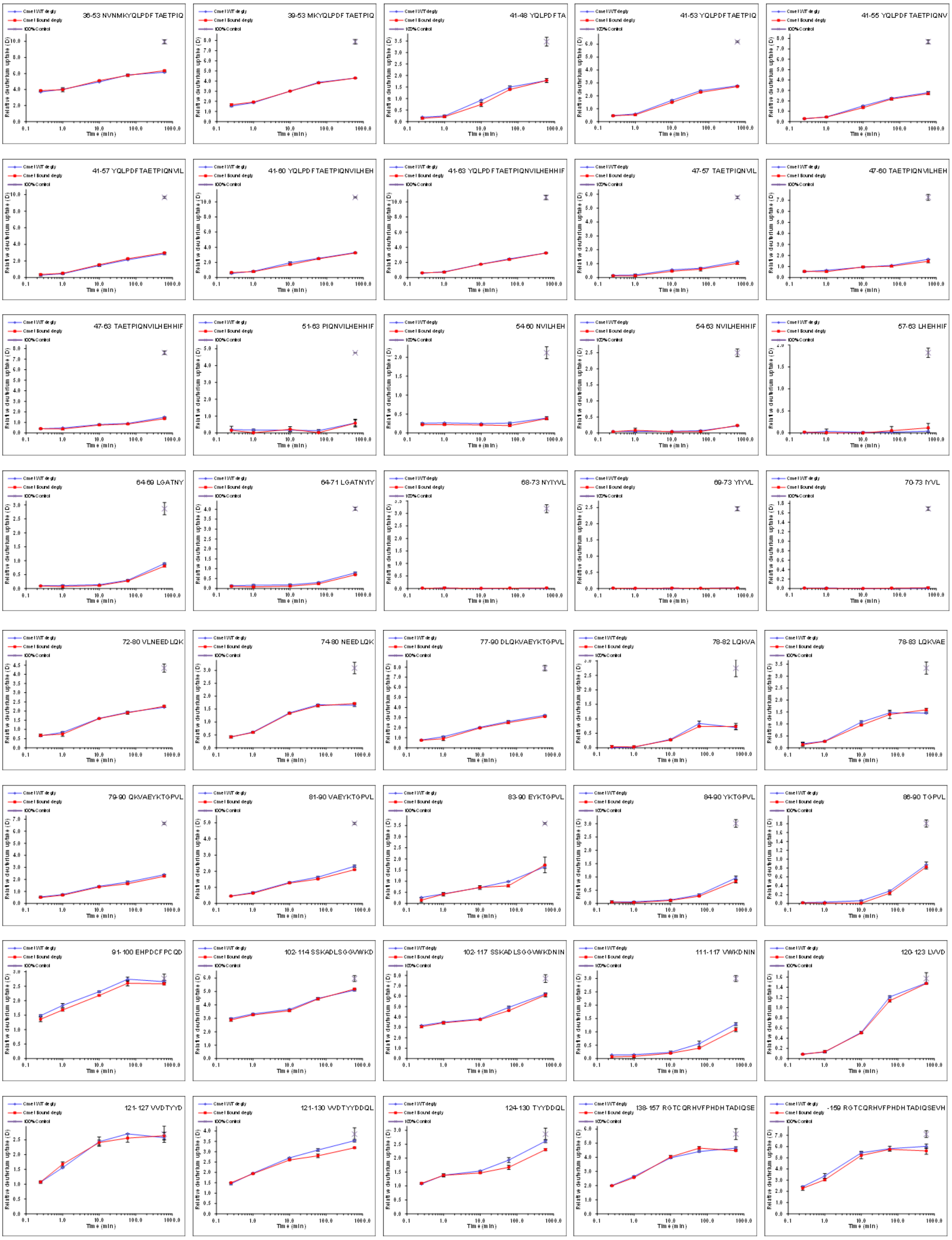

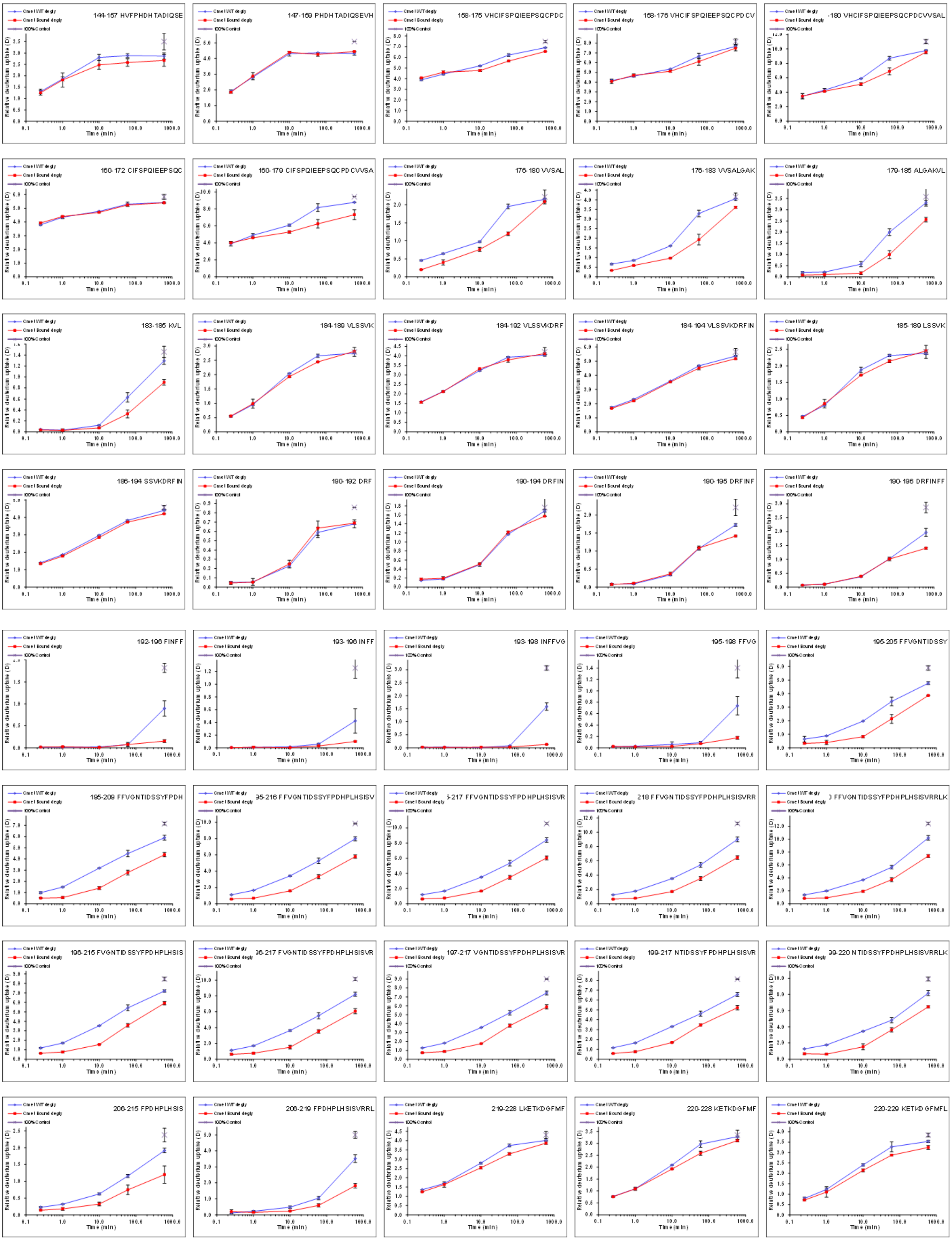
144-157 HVFPHDHTADIQSE
147-159 PHDHTADIQSEVH
158-175 VHCIFSPQIEEPSQCPDC
158-176 VHCIFSPQIEEPSQCPDCV
160-172 CIFSPQIEEPSQC
160-179 CIFSPQIEEPSQCPDCVVSA
176-180 VVSAL
176-183 VVSALGAK
179-185 ALGAKVL
183-185 KVL
184-189 VLSSVK
184-192 VLSSVKDRF
184-194 VLSSVKDRFIN
185-189 LSSVK
186-194 SSVKDRFIN
190-192 DRF
190-194 DRFIN
190-195 DRFINF
190-196 DRFINFF
192-196 FINFF
193-196 INFF
193-198 INFFVG
195-198 FFVG
195-205 FFVGNTIDSSY
195-209 FFVGNTIDSSYFPDH
206-215 FPDHPLHSIS
206-219 FPDHPLHSISVRRL
219-228 LKETKDGFMF
220-228 KETKDGFMF
220-229 KETKDGFMFL
Time (min)
Relative deuterium uptake (D)
100% Control

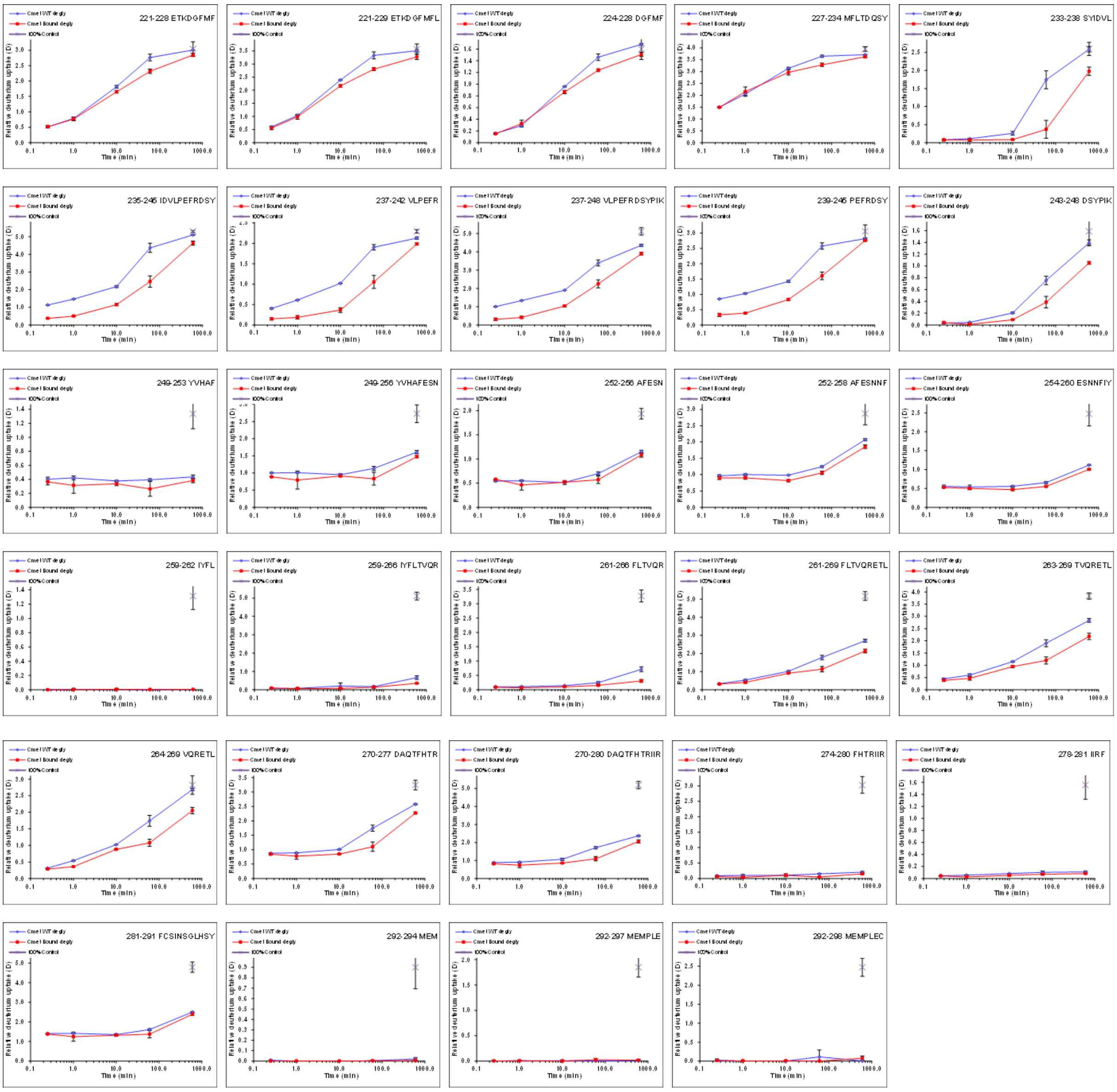
221-228 ETKDGFMF
221-229 ETKDGFMFL
224-228 DGFMF
227-234 MFLTDQSY
233-238 SYIDVL
235-246 IDVLPEFRDSY
237-242 VLPEFR
237-248 VLPEFRDSYPIK
239-246 PEFRDSY
243-248 DSYPIK
249-253 YVHAF
249-256 YVHAFESN
252-256 AFESN
252-258 AFESNNF
254-260 ESNNFIY
259-262 IYFL
259-266 IYFLTVQR
261-266 FLTVQR
261-269 FLTVQRETL
263-269 TVQRETL
264-269 VQRETL
270-277 DAQTFHTR
270-280 DAQTFHTRIIR
274-280 FHTRIIR
278-281 IIRF
281-291 FCSINSGLHSY
292-294 MEM
292-297 MEMPLE
292-298 MEMPLEC
Relative deuterium uptake (D)
Time (min)
100% Control

#### C-met β Chain

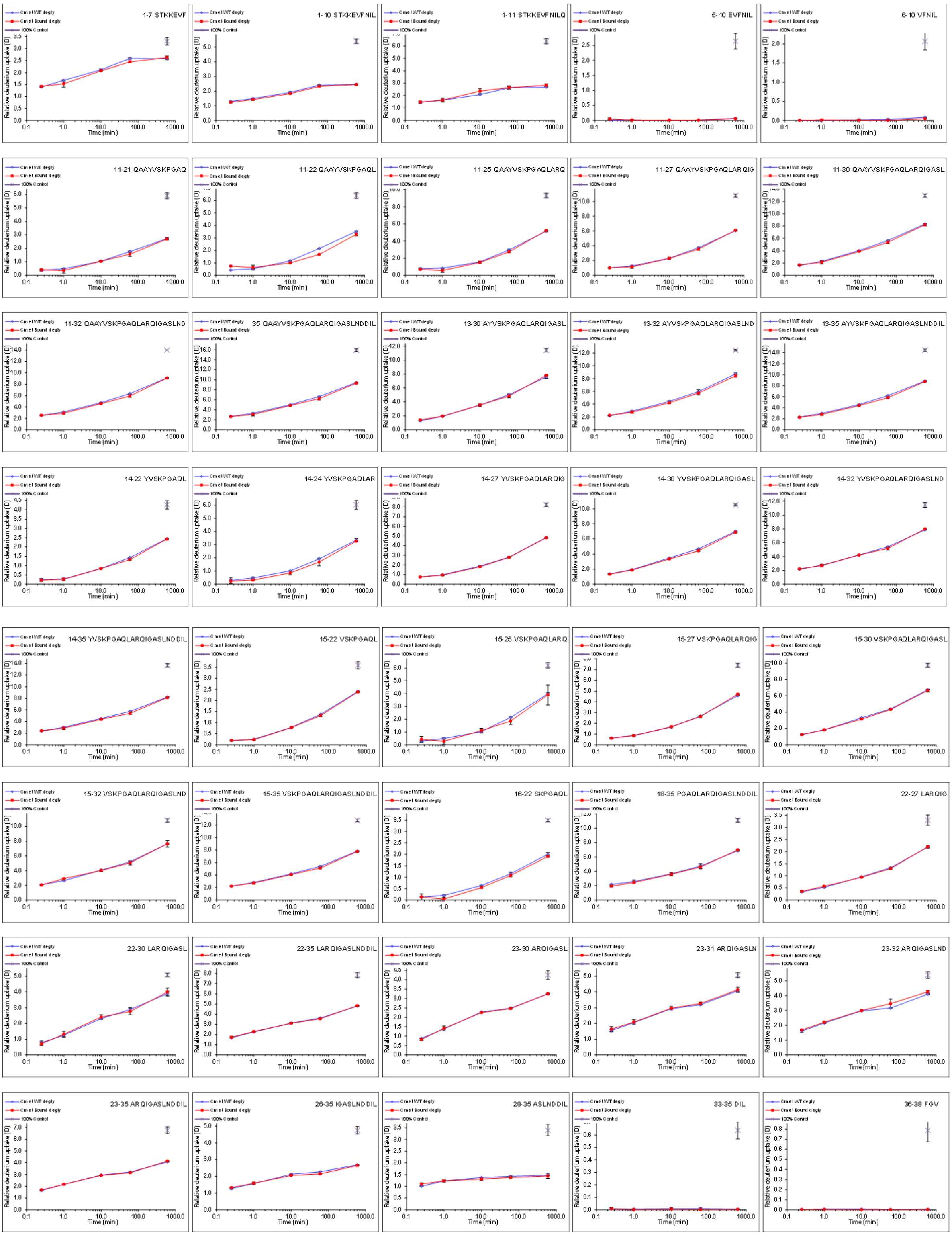
1-7 STKKEVF
1-10 STKKEVFNIL
1-11 STKKEVFNILQ
5-10 EVFNIL
6-10 VFNIL
11-21 QAAYVSKPGAQ
11-22 QAAYVSKPGAQL
11-25 QAAYVSKPGAQLARQ
11-27 QAAYVSKPGAQLARQIG
11-30 QAAYVSKPGAQLARQIGASL
11-32 QAAYVSKPGAQLARQIGASLND
35 QAAYVSKPGAQLARQIGASLNDDIL
13-30 AYVSKPGAQLARQIGASL
13-32 AYVSKPGAQLARQIGASLND
13-35 AYVSKPGAQLARQIGASLNDDIL
14-22 YVSKPGAQL
14-24 YVSKPGAQLAR
14-27 YVSKPGAQLARQIG
14-30 YVSKPGAQLARQIGASL
14-32 YVSKPGAQLARQIGASLND
14-35 YVSKPGAQLARQIGASLNDDIL
15-22 VSKPGAQL
15-25 VSKPGAQLARQ
15-27 VSKPGAQLARQIG
15-30 VSKPGAQLARQIGASL
15-32 VSKPGAQLARQIGASLND
15-35 VSKPGAQLARQIGASLNDDIL
16-22 SKPGAQL
18-35 PGAQLARQIGASLNDDIL
22-27 LARQIG
22-30 LARQIGASL
22-35 LARQIGASLNDDIL
23-30 ARQIGASL
23-31 ARQIGASLN
23-32 ARQIGASLND
23-35 ARQIGASLNDDIL
26-35 IGASLNDDIL
28-35 ASLNDDIL
33-35 DIL
36-38 FGV
Time (min)
Relative deuterium uptake (D)
100% Control

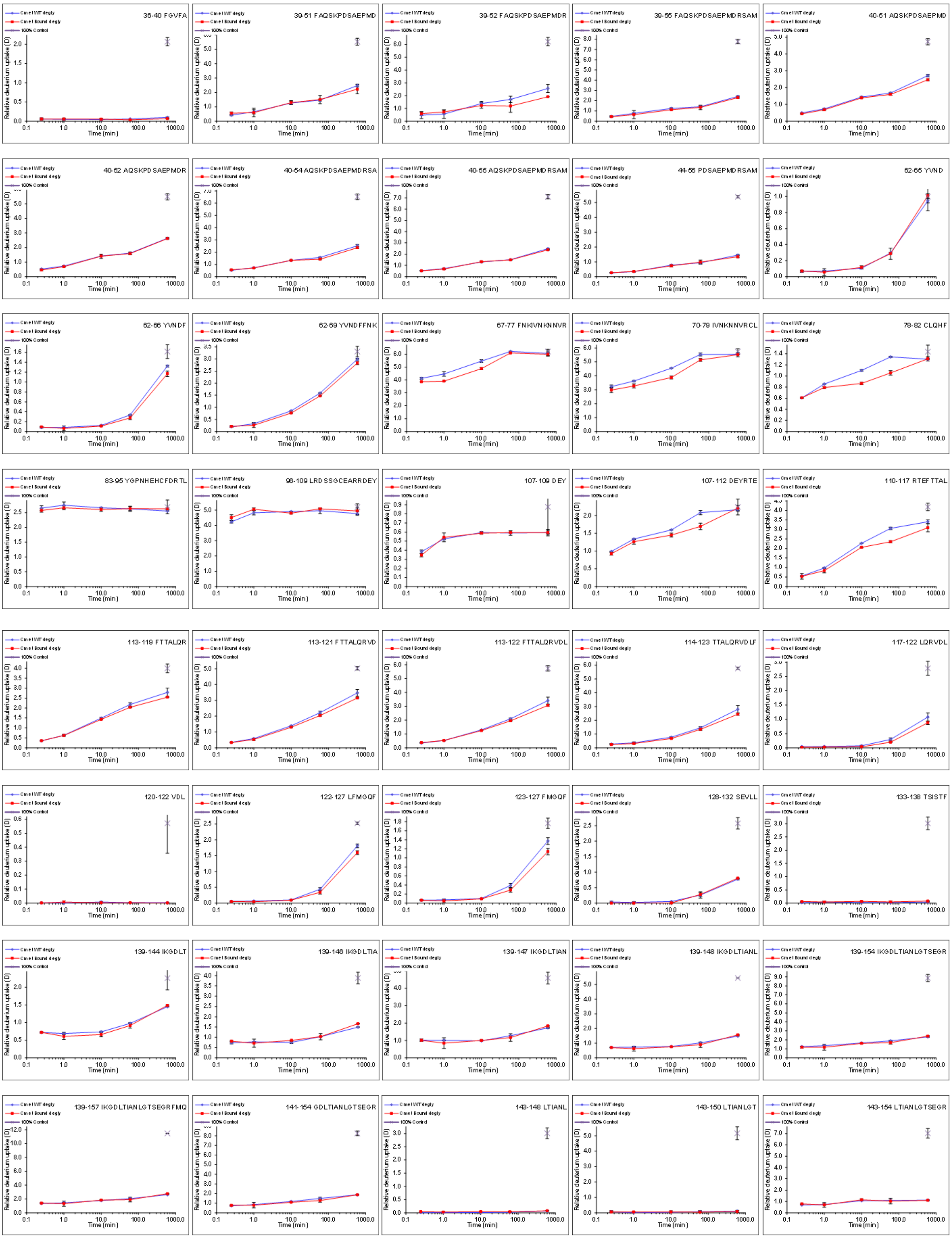
36-40 FGVFA
39-51 FAQSKPDSAEPMD
39-52 FAQSKPDSAEPMDR
39-55 FAQSKPDSAEPMDRSAM
40-51 AQSKPDSAEPMD
40-52 AQSKPDSAEPMDR
40-54 AQSKPDSAEPMDRSA
40-55 AQSKPDSAEPMDRSAM
44-55 PDSAEPMDRSAM
62-65 YVND
62-66 YVNDF
62-69 YVNDFFNK
67-77 FNKIVNKNNVR
70-79 IVNKNNVRCL
78-82 CLQHF
83-95 YGPNHEHCFDRTL
96-109 LRDSSGCEARRDEY
107-109 DEY
107-112 DEYRTE
110-117 RTEFTTAL
113-119 FTTALQR
113-121 FTTALQRVD
113-122 FTTALQRVDL
114-123 TTALQRVDLF
117-122 LQRVDL
120-122 VDL
122-127 LFMGQF
123-127 FMGQF
128-132 SEVLL
133-138 TSISTF
139-144 IKGDLT
139-146 IKGDLTIA
139-147 IKGDLTIAN
139-148 IKGDLTIANL
139-154 IKGDLTIANLGTSEGR
139-157 IKGDLTIANLGTSEGRFMQ
141-154 GDLTIANLGTSEGR
143-148 LTIANL
143-150 LTIANLGT
143-154 LTIANLGTSEGR
Relative deuterium uptake (D)
Time (min)

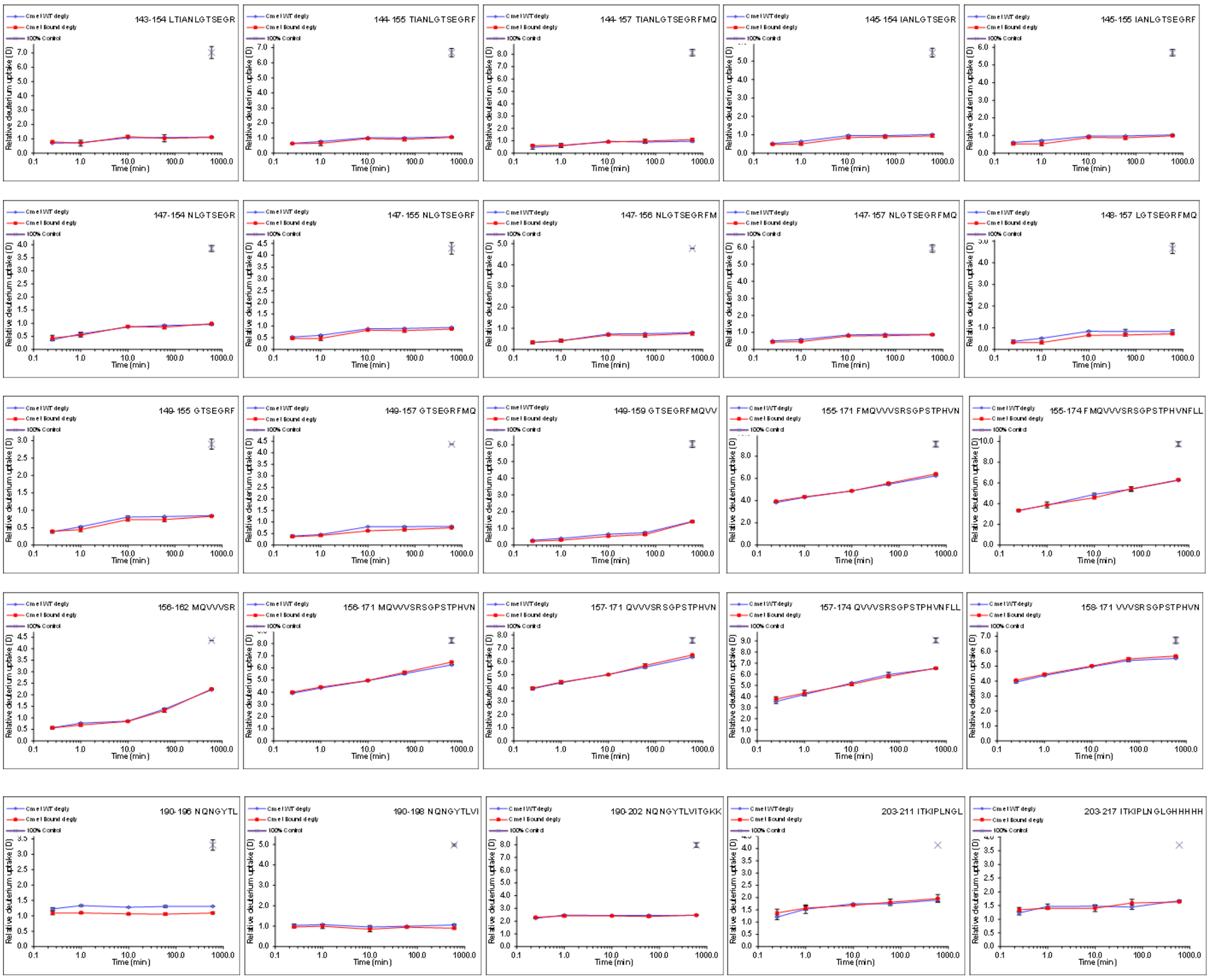

*Figure S11: HDX uptake plots of peptides from the C-Met epitope mapping. The HDX of C-Met with mAb is highlighted red, C-met without mAb is highlighted blue and maximally-labelled control samples is shown as a cross. Error bars represent the standard deviation from three technical replicate measurements.*

### 7 VISTA

#### 7.1 HDX-LC-MS experimental set-up with on-line pepsin digestion and deglycosylation

HDX samples were analyzed on an LC-MS system containing an RSLCnano system (UltiMate 3000 RSLCnano, Thermo Fisher Scientific) with two pumps, a chilling device for chromatography (MéCour Temperature Control, USA) and a mass spectrometer (Orbitrap Eclipse Tribrid, Thermo Fisher Scientific).

The chilling device (17 x 23 x 23 cm, HxWxL) was temperature-controlled by constant flow of propylene glycol (40%) chilled to -4 °C through tubes inside the wall of the chamber using a MiniChiller 280 pump (Huber, Offenburg; Germany). Inside the chilling device following components were kept at 0 °C: an eluent-cooling loop (stainless steel capillary; 0.1 x 150 mm), a 2-way-6-port valve (7725i, Rheodyne) for column switching, a manual injector valve (2-way-6-port valve, Rheodyne) with a 20 µL stainless-steel sample loop (Rheodyne), a pepsin-digestion column (2.1 x 30 mm, Enzymate$^{TM}$ BEH Pepsin Column, Waters, Germany), a PNGase Rc-deglycosylation column (2.1 x 30 mm with PNGase Rc-POROS$^{TM}$ 20 AL beads packed by Dr. Maisch HPLC GmbH.), a trapping column (Acquity BEH VanGuard C18 pre-column, 2.1 x 5 mm, 1.7 µm, 130 Å, Waters, Germany), a microLC column (ACQUITY BEH C18, 1.7 µm, 300 Å, 1 mm x 50 mm (Waters GmbH, Germany)) and a waste bottle.

The loading pump of the RSLCnano system was used for on-line pepsin digestion and for subsequent on-line deglycosylation. Both columns were placed one after another. Eluent was water with 0.1 % formic acid (pH 2.5) at a flow rate of 50 µL/min for 2.5 minutes to account for on-column pepsin digestion time. From 2.5 to 3 minutes the flow was increased to 100 µL. The flow was than kept at 100 µL/min for 5 minutes to account for on-line deglycosylation and trapping of the peptides on the VanGuard column. After 5 minutes, the valve was switched, and the trapped peptides were transferred to the analytical column using the Nano/Cap System (NCS) pump of the RSLC system. A linear 10-min gradient with a flow rate of 50 µl/min from 10 % B to 60 % B followed by column washing steps was applied. Solvent A was 0.1% (v/v) formic acid and solvent B was 80% acetonitrile (v/v) with 0.1% formic acid (v/v). After each analytical run, a column-wash gradient was performed.

MS and MSMS analysis were performed with 60,000 resolution using the orbitrap and the ion trap with data dependent HCD fragmentation and rapid activation of the Orbitrap Eclipse Tribrid instrument, respectively. Ion source and detection parameters were used as follows: m/z: 300-2000; positive ion spray voltage and an ion transfer tube temperature of 275°C, sheath gas flow rate of 25; aux gas flow rate of 5; S-lens RF level of 50, spray voltage of 3.5 kV and a capillary temperature of 300°C.

#### 7.2 HDX uptake plots of VISTA

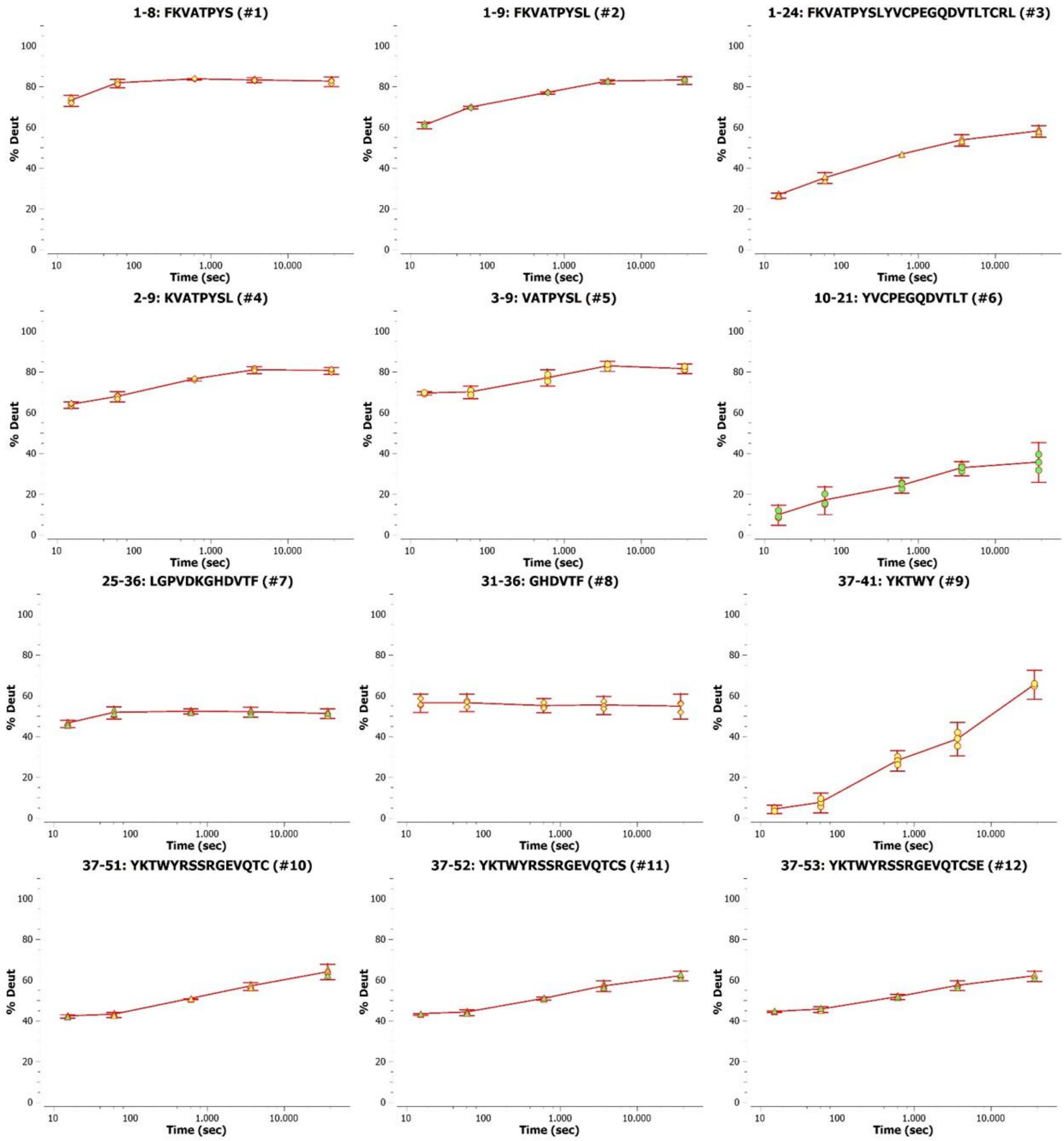

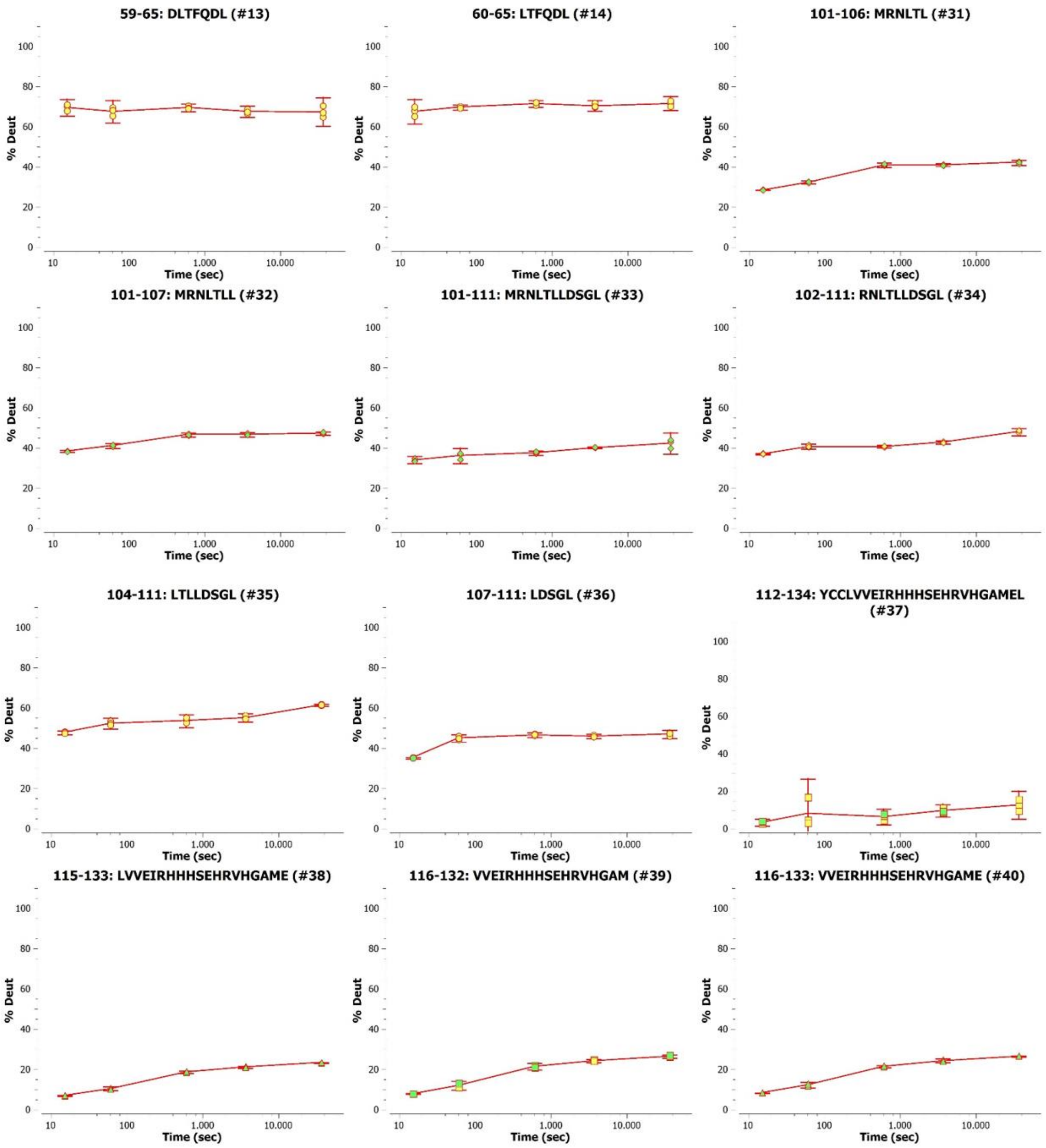
59-65: DLTFQDL (#13)
60-65: LTFQDL (#14)
101-106: MRNLTL (#31)
101-107: MRNLTLL (#32)
101-111: MRNLTLLDSGL (#33)
102-111: RNLTLLDSGL (#34)
104-111: LTLLDSGL (#35)
107-111: LDSGL (#36)
112-134: YCCLVVEIRHHHSEHRVHGAMEL (#37)
115-133: LVVEIRHHHSEHRVHGAME (#38)
116-132: VVEIRHHHSEHRVHGAM (#39)
116-133: VVEIRHHHSEHRVHGAME (#40)
% Deut
Time (sec)
0
20
40
60
80
100
10
100
1.000
10.000

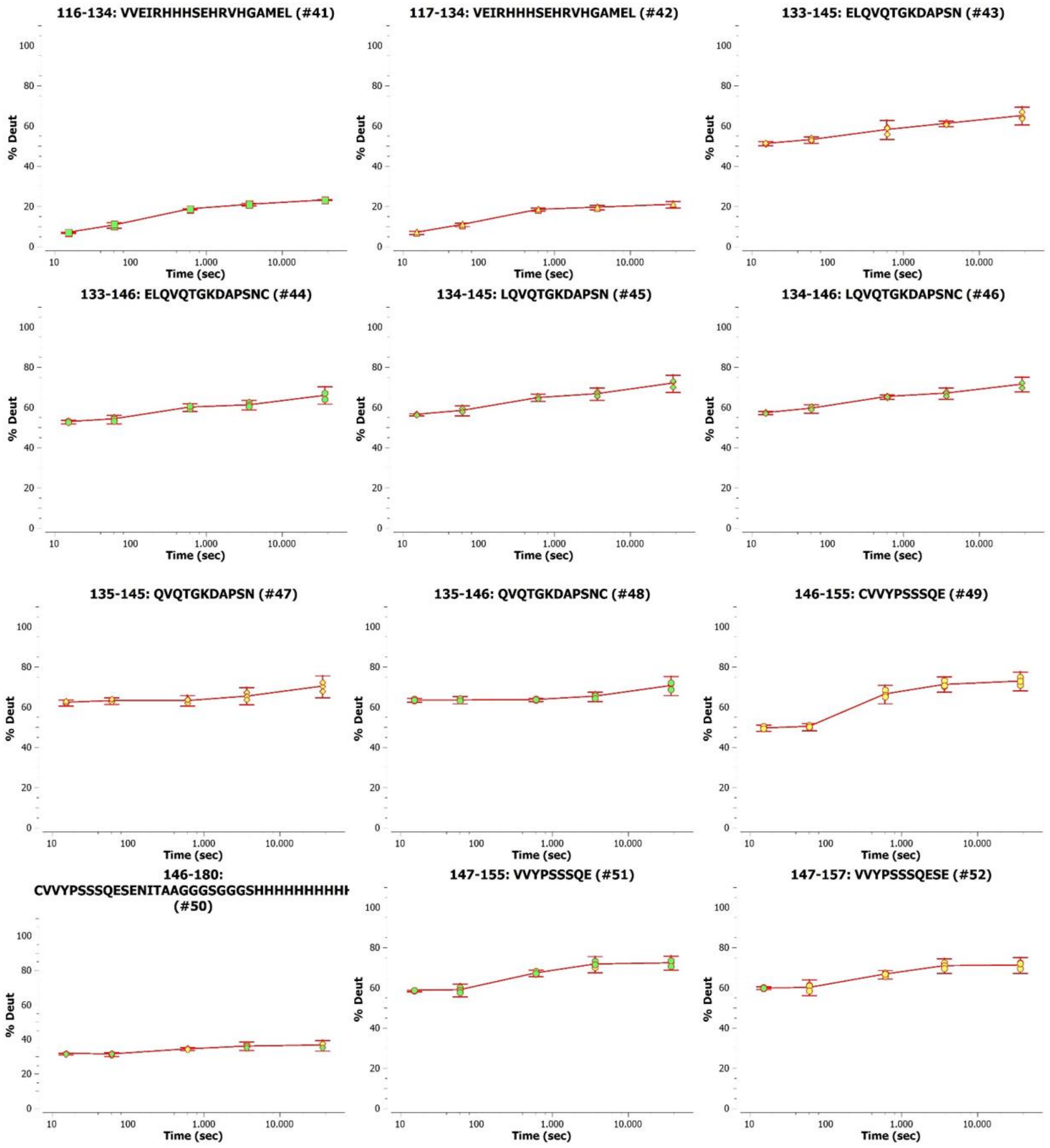
116-134: VVEIRHHHSEHRVHGAMEL (#41)
117-134: VEIRHHHSEHRVHGAMEL (#42)
133-145: ELQVQTGKDAPSN (#43)
133-146: ELQVQTGKDAPSNC (#44)
134-145: LQVQTGKDAPSN (#45)
134-146: LQVQTGKDAPSNC (#46)
135-145: QVQTGKDAPSN (#47)
135-146: QVQTGKDAPSNC (#48)
146-155: CVVYPSSSQE (#49)
146-180: CVVYPSSSQESENITAAGGGSGGGSHHHHHHHHHH (#50)
147-155: VVYPSSSQE (#51)
147-157: VVYPSSSQESE (#52)
% Deut
0
20
40
60
80
100
Time (sec)
10
100
1,000
10,000

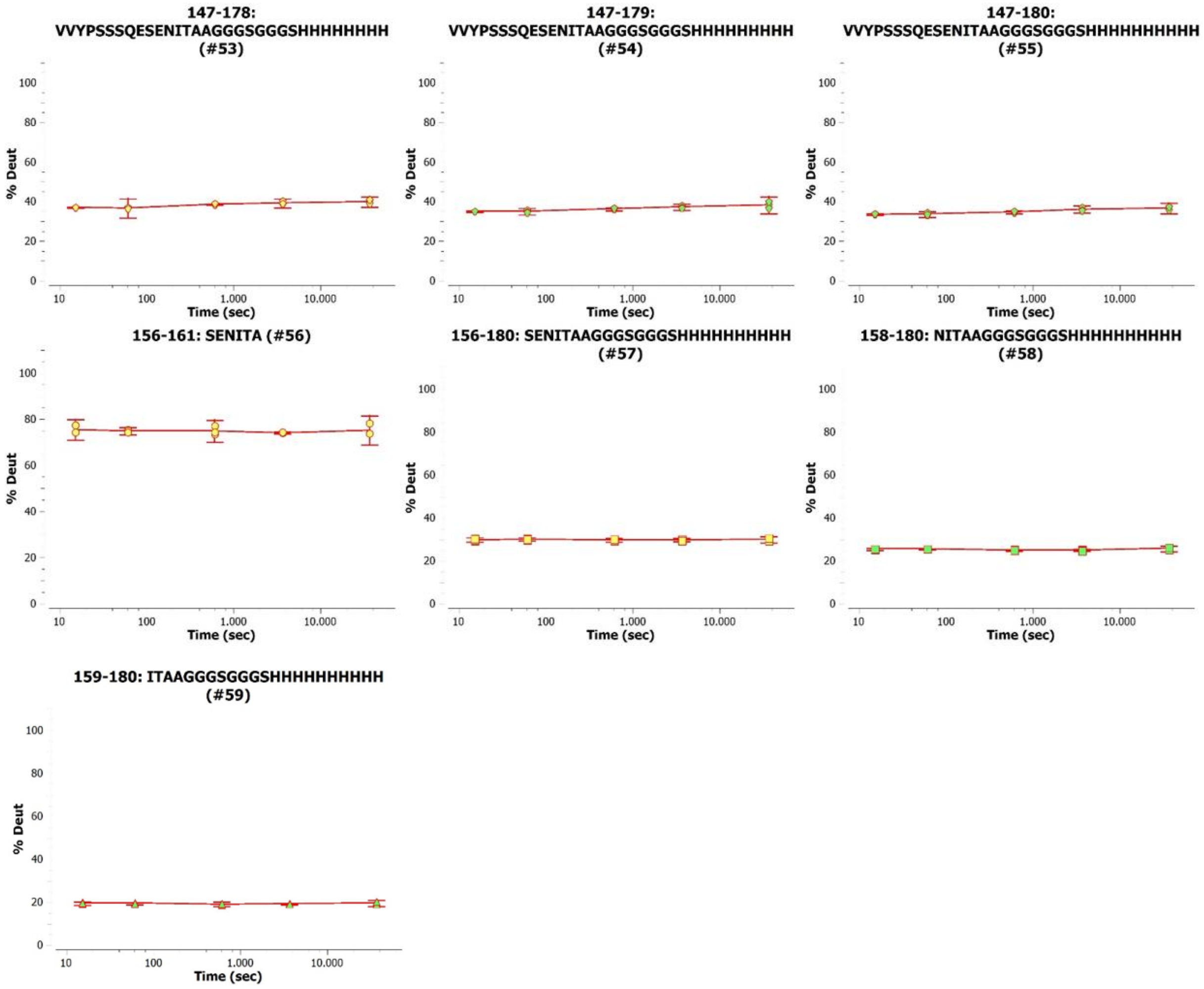

*Figure S12: HDX uptake plots of peptic peptides of VISTA. Deuterium uptake of each peptide is normalized to the exchangeable amino acid residues (number of amino acids minus the first 2 N –terminal residues and proline). Error bars show the significance interval on 95% confidence.*
